# Prokrustean Graph: A substring index for rapid k-mer size analysis

**DOI:** 10.1101/2023.11.21.568151

**Authors:** Adam Park, David Koslicki

## Abstract

The widespread adoption of *k*-mers in bioinformatics has led to efficient methods utilizing genomic sequences in a variety of biological tasks. However, understanding the influence of *k*-mer sizes within these methods remains a persistent challenge, as the outputs of complex bioinformatics pipelines obscure this influence with various noisy factors. The choice of *k*-mer size is often arbitrary, with justification frequently omitted in the literature and method tutorials. Furthermore, recent methods employing multiple *k*-mer sizes encounter significant computational challenges.

Nevertheless, most methods are built on well-defined objects related to *k*-mers, such as de Bruijn graphs, Jaccard similarity, Bray-Curtis dissimilarity, and *k*-mer spectra. The role of *k*-mer sizes within these objects is more intuitive and can be described by numerous quantities and metrics. Therefore, exploring these objects across *k*-mer sizes opens opportunities for robust analyses and new applications. However, the evolution of *k*-mer objects with respect to *k*-mer sizes is surprisingly elusive.

We introduce a novel substring index, the Pro*k*rustean graph, that elucidates the transformation of *k*-mer sets across *k*-mer sizes. Our framework built upon this index rapidly computes *k*-mer-based quantities for all *k*-mer sizes, with computational complexity independent of the size range and dependent only on maximal repeats. For example, counting maximal simple paths in de Bruijn graphs for *k* = 1, …, 100 is achieved in seconds using our index on a gigabase-scale dataset. We present a variety of such experiments relevant to pangenomics and metagenomics.

The Pro*k*rustean graph is space-efficiently constructed from the Burrows-Wheeler Transform. Through this construction, it becomes evident that other modern substring indices inherently face difficulties in exploring *k*-mer objects across sizes, which motivated our data structure.

Our implementation is available at: https://github.com/KoslickiLab/prokrustean.

## 1 Introduction

As the volume and number of sequencing reads and reference genomes grow every year, *k*-mer-based methods continue to gain popularity in computational biology. This simple approach—cutting sequences into substrings of a fixed length—offers significant benefits across various disciplines. Biologists consider *k*-mers as intuitive markers representing biologically significant patterns, bioinformaticians easily formulate novel methods by regarding *k*-mers as fundamental units that reflect their original sequences, and engineers leverage their fixed-length nature to optimize computational processes at a very low level. However, understanding the most crucial parameter, *k*, remains elusive, thereby leaving the accuracy of experimental results obscure to practitioners.

Two central challenges emerge in *k*-mer-based methods. First, the selection of the *k*-mer size is often arbitrary, despite its well-recognized influence on outcomes. This issue, though widely acknowledged, remains insufficiently addressed in the literature, with little formal guidance on determine an optimal *k* size for applications. The reasoning behind these choices is frequently unclear, typically confined to specific, unpublished experimental analyses (e.g., “we found that *k*=31 was appropriate… “). Second, methods attempting to utilize multiple *k*-mer sizes encounter significant computational burdens. The accuracy benefits of incorporating multiple *k*-mer sizes are overshadowed by the escalating computational costs with each additional *k*-mer size. Consequently, researchers often resort to “folklore” *k* values based on prior empirical results.

The influence of *k*-mer sizes within each method is a complex function reflecting bioinformatics pipelines that process *k*-mers. As biological adjustments and engineering strategies complicate the pipelines, their outputs obscure the impact of *k*-mer sizes with noisy factors, as simplified in the following abstraction:

> pipeline (`k-mer-based objects(sequences, k)`, biological adjustments, engineering).

Despite the complexity of the pipelines, there always exist mathematically well-defined *k*-mer-based objects that form the foundation of method formulation. These objects abstract the utilization of *k*-mers and depend solely on sequences and *k*-mer sizes. Thus, they offer potential for generalizability and quantification of the influence of *k*-mers in methods. Indeed, intuitive quantities have been derived from *k*-mer-based objects; however, there are challenges in actually computing them, as detailed in several examples we now present.

In genome analysis, the number of distinct *k*-mers is often used to reflect the complexity of genomes. Although the number varies by *k*-mer sizes, large data sizes restrict experiments to a few *k* values [9, 15, 39]. A recent study suggests that the number of distinct *k*-mers across all *k*-mer sizes provides additional insights into pangenome complexity [10]. Furthermore, the frequencies of *k*-mers provide richer information, as discussed in [1, 46], but their computation becomes more complicated and sometimes impractical, even with a fixed *k*-mer size [7].

In comparative analyses, Jaccard Similarity is utilized for genome indexing and searching by being approximated through hashing techniques [26, 34]. Experiments attempting to assess the influence of *k*-mer sizes on Jaccard Similarity must undergo tedious iterations through various *k*-mer sizes [8]. Additionally, Bray-Curtis dissimilarity is a *k*-mer frequency-based measure frequently used to compare metagenomic samples, where experimentalists face resource limitations due to the growing size of sequencing data, even with a fixed *k*-mer size. Yet, the demand for analyzing multiple *k*-mer sizes continues to increase [23, 35, 36].

Genome assemblers utilizing *k*-mer-based de Bruijn graphs are probably the most sensitive to *k*-mer sizes, and those employing multi-*k* approaches face significant computational challenges. The choice of *k*-mer sizes is particularly crucial in *de novo* assemblers, yet they predominantly rely on heuristic methods [19, 27, 38]. The topological features of de Bruijn graphs are succinctly summarized by the compacting process, where vertices represent simple paths called unitigs, but exploring these features across varying *k*-mer sizes is computationally intensive. Furthermore, the development of multi-*k* de Bruijn graphs continues to be a challenging and largely theoretical endeavor [22, 40, 43], with no substantial advancements following the heuristic selection of multiple *k*-mer sizes implemented by metaSPAdes [33].

These examples motivated the pressing need to generalize the exploration of *k*-mer-based objects across varying *k*-mer sizes. We introduce a computational framework for rapidly computing quantities derived from *k*-mer-based objects across a range of *k* sizes. The objective is clarified in Section 2, with experimental results in Section 3. The underlying substring index and its usage are briefly presented in Section 4 and Section 5.

Due to the length of proofs and algorithms, construction algorithms are left in Supplementary materials. We emphasize that distinctions between the new substring index and other modern substring indexes are only briefly covered in Section 5, while the construction algorithms provide essential insights. The new index can be built from the Burrows-Wheeler Transform (BWT), but the BWT alone is unlikely to achieve the same efficiency as our framework. It appears that modern substring indexes based on longest common prefixes, such as the BWT, suffix tree, FM-index, and r-index, share similar limitations.

## 2 Setup and main theorem

Let *Σ* be our alphabet and let *S* ∈*Σ*^∗^ represent a finite-length string, and 𝒰⊂*Σ*^∗^ represent a set of strings. Consider the following preliminary definitions and main result detailing the Pro*k*rustean graph utility:

### Definition 1.

– *A <u>region</u> in the string S is a triple* (*S, i, j*) *such that* 1 ≤ *i* ≤ *j* ≤|*S*|. *\*\* See the example after definition 2 which justifies this non-standard region notation* (*S, i, j*).
– *The <u>string</u> of a region is the corresponding substring: str*(*S, i, j*) := *S*_*i*_*S*_*i*+1_…*S*_*j*_.
– *The <u>size</u> or <u>length</u> of a region is the length of its string:* | (*S, i, j*) | := *j* −*i* + 1.
– *An <u>extension</u> of a region* (*S, i, j*) *is a larger region* (*S, i*^*′*^, *j*^*′*^) *including* (*S, i, j*):

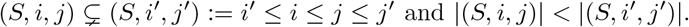
– *The <u>occurrences</u> of S in 𝒰 are those regions in strings of 𝒰 corresponding to S:*

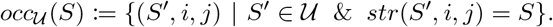
– *S is a <u>maximal repeat</u> in 𝒰 if it occurs at least twice and more than any of its extensions:* |*occ*_*𝒰*_ (*S*)| *>* 1, *and for all S*^*′*^ *a superstring of S*, |*occ*_*𝒰*_ (*S*)| *>* |*occ*_*𝒰*_ (*S*^*′*^)|. *I*.*e*., *a maximal repeat S is one that occurs more than once in* 𝒰 *(i*.*e. is a repeat), and any extension of which has a lower frequency in* 𝒰 *than S*.
– *Let* ℛ _𝒰_ *represent the set of all maximal repeats in* 𝒰.

### Theorem 1. (Main Result)

*Given a set of sequences* 𝒰, *there exists a substring representation* 𝒢 _𝒰_ *of size O*(|*Σ*|| ℛ _𝒰_ |). 𝒢 _𝒰_ *can be used to compute <u>k-mer quantities</u> of* 𝒰 *for all k* = 1, …, *k*_max_ *within O*(|𝒢 _𝒰_ |) *time and space, where k*_max_ *is the length of the longest sequence in* 𝒰. *The <u>k-mer quantities</u> are defined as follows:*

– *(Counts) The number of distinct k-mers, as used in genome cardinality DandD [10]*.
– *(Frequencies) k-mer Bray-Curtis dissimilarities, as used in comparing metagenomic samples [36]*.
– *(Extensions) The number of maximal unitigs in k-mer de Bruijn graphs, as used in analyses of reference genomes and genome assembly [29, 42]*.
– *(Occurrences) The number of edges with annotated length of k in an overlap graph. [45]*.

*Furthermore*, 𝒢 _𝒰_ *can be constructed in O*(|*bwt*|+| 𝒢 _𝒰_ |) *time and space utilizing bwt, a BWT representation of* 𝒰.

*Proof*. Section 4 defines 𝒢 _𝒰_ and derives its size *O*(|*Σ*|| ℛ _𝒰_ |). Section 5 introduces the computation of *k*-mer quantities, and the construction algorithm is introduced in the Supplementary Section S3. □

𝒢 _*𝒰*_ is termed the Pro*k*rustean graph of 𝒰. The main contribution is that the time complexity for computing *k*-mer quantities for all *k* ∈ [1, *k*_max_] is independent of the range, specifically *O*(|𝒢 _𝒰_ |). Notably, |𝒢 _𝒰_| is sublinear relative to *N*, where *N* is the cumulative sequence length of. In contrast, any direct computation of *k*-mer quantities from 𝒰 for a fixed *k*-mer size demands, at best, *O*(*N*) time. Extending this computation naively to *k* = 1, …, *k*_max_ increases the complexity to *O*(*N* ^2^) to cover all possible substrings.

## 3 Results

The construction and four applications of the Pro*k*rustean graph were conducted with various datasets that were randomly selected to represent a broad range of sizes, sequencing technologies, and biological origins. Diverse sequencing datasets were collected, including two metagenome short-read datasets, one metagenome long-read dataset, and two human transcriptome short-read datasets. The full list of datasets is provided in section S1. All experiments were performed on a Mac Studio equipped with an Apple M1 Max of 10 CPU cores and 64 GB memory.

### 3.1 Pro*k*rustean graph construction with k_*min*_

The construction algorithm is introduced in the Supplementary Section S3. The Pro*k*rustean graph grows with the number of maximal repeats, i.e., *O*(|𝒢 _𝒰_ |) = *O*(|*Σ*| *·* |ℛ_𝒰_ |). Also, the size of 𝒢 _𝒰_ can be controlled by dropping maximal repeats of length below *k*_*min*_. Then 𝒢_*𝒰*_ requires *O*(|*Σ*| *·* |ℛ_𝒰_ (*k*_*min*_)|) space where ℛ_𝒰_ (*k*_*min*_) is the set of maximal repeats of length at least *k*_*min*_. Then, subsequent sections cover the results of four applications that compute *k-mer quantities* for *k* = *k*_*min*_, …, *k*_*max*_. We assume *k*_*max*_ is always less than |𝒢 _𝒰_|, even if the largest possible value is chosen, which is typical for sequencing data. Figure 1 shows how the graph size (*O*(|*Σ*| *·* |ℛ_*𝒰*_ (*k*_*min*_)|)) of short sequencing reads decreases as *k*_*min*_ increases.

**Fig. 1:**
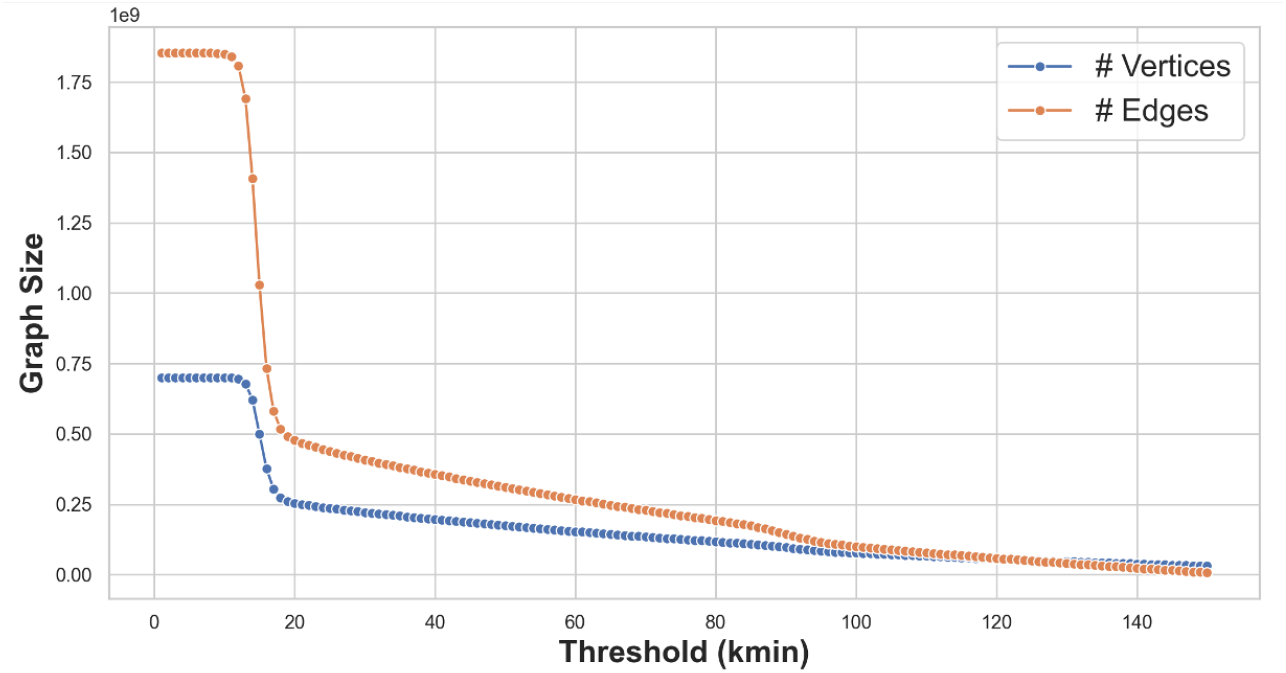
Size decay of a Pro*k*rustean graph as a function of *k*_*min*_. The sizes of Pro*k*rustean graphs, i.e., number of vertices and edges, versus *k*_*min*_ values for a short read dataset ERR3450203 are depicted. The size of the graph rapidly drops around *k*_*min*_ = [11, …, 19], meaning the maximal repeats and locally-maximal repeat regions are dense when *k*_*min*_ is under 11. The number of edges falls again around *k*_*min*_ = 100 because the graph eventually becomes fully disconnected.

Table 1 shows that the size of the Pro*k*rustean graphs is sublinear to that of their inputs. Also, the space occupancy is close to the inputs (read files) with practical *k*_*min*_ values. We used *k*_*min*_ = 30 that is common in genome assemblers and comparative microbiome analyses.

**Table 1:** Pro*k*rustean graph construction introduced in Section S3. Symbols M, B, mb, and gb denote millions, billions, megabytes, and gigabytes, respectively. *N* represents the cumulative sequence length of 𝒰. 𝒱 _𝒰_ is the vertex set and ℰ _𝒰_ is the edge set of 𝒢 _𝒰_. Since | ℰ _𝒰_ | ≪ *N*, the time complexity of subsequent applications *O*(|𝒢_𝒰_ |) := *O*(|ℰ _𝒰_ |) is sublinear to the input, even if all *k*-mer sizes are considered.

| dataset | reads | N | $k_{min}$ | time | memory | output size | $\vert \mathcal{V}_{\mathcal{U}}\vert $ | $\vert \mathcal{E}_{\mathcal{U}}\vert $ |
| --- | --- | --- | --- | --- | --- | --- | --- | --- |
| SRR20044276 (metagenomic) | 0.94M | 72M | 1 | 20s | 200mb | 40mb | 8M | 21M |
| SRR20044276 (metagenomic) | 0.94M | 72M | 20 | 13s | 180mb | 35mb | 2.7M | 5.4M |
| ERR3450203 (metagenomic) | 55M | 4.5B | 20 | 29min | 12.7gb | 3.1gb | 254M | 477M |
| ERR3450203 (metagenomic) | 55M | 4.5B | 30 | 28min | 11gb | 2.7gb | 221M | 407M |
| SRR18495451 (metagenomic/long) | 0.83M | 11.3B | 30 | 90min | 43gb | 14gb | 890M | 1.7B |
| SRR7130905 (human/rna-seq) | 45M | 6.8B | 30 | 39min | 18gb | 4gb | 280M | 580M |
| SRR21862404 (human/rna-seq) | 336M | 22.3B | 30 | 99min | 37gb | 5.9gb | 330M | 800M |

Regarding subsequent results, note that, to our knowledge, no existing computational techniques perform the exact same tasks across a range of *k*-mer sizes, making performance comparisons inherently “unfair.” Readers are encouraged to focus on the overall time scale to gauge the efficiency of the Pro*k*rustean graph. Additionally, discrepancies in output comparisons may arise due to practice-specific configurations in computational tools, such as the use of only ACGT (i.e., no “N”) and canonical *k*-mers in KMC. Consequently, we tested correctness on GitHub^1^ using our brute-force implementations.

### 3.2 Counting distinct *k*-mers for *k* = *k*_*min*_, …, *k*_*max*_

The number of distinct *k*-mers is commonly used to assess the complexity and structure of pangenomes and metagenomes [13, 44]. A recent study by [10] spans multiple *k* contexts, suggesting that the number of *k*-mers across all *k*-mer sizes should be used to estimate genome sizes. The Supplementary Algorithm S1 counts distinct *k*-mers for all *k*-mer sizes in time *O*(| 𝒢 _𝒰_ |). We employed KMC [28], an optimized *k*-mer counting library designed for a fixed *k* size, for comparison, and used it iteratively for all *k* values.

### 3.2 Counting simple paths in de Bruijn graphs for *k*_*min*_, …, *k*_*max*_

A maximal unitig of a de Bruijn graph represents a simple path that cannot be extended further. Maximal unitigs are vertices of a compacted de Bruijn graph [20], reflecting a topological characteristic of the graph. More maximal unitigs generally indicate more complex graph structures, which significantly impact genome assembly performance [3, 14]. Therefore, their number across multiple *k*-mer sizes reflects the influence of *k*-mer sizes on the complexity of assembly. Algorithm S3 counts maximal unitigs for all *k*-mer sizes in time *O*(|𝒢 _𝒰_ |).

We utilized GGCAT [20] for comparisons, a mature library constructing compacted de Bruijn graphs of a fixed *k*-mer size. We assumed their construction is equivalent to counting maximal unitigs. Again, computing maximal unitigs with the Pro*k*rustean graph becomes more efficient as additional *k*-mer sizes are considered. Table 3 displays the comparison, and Figure 2 illustrates one of the outputs.

**Table 2:**
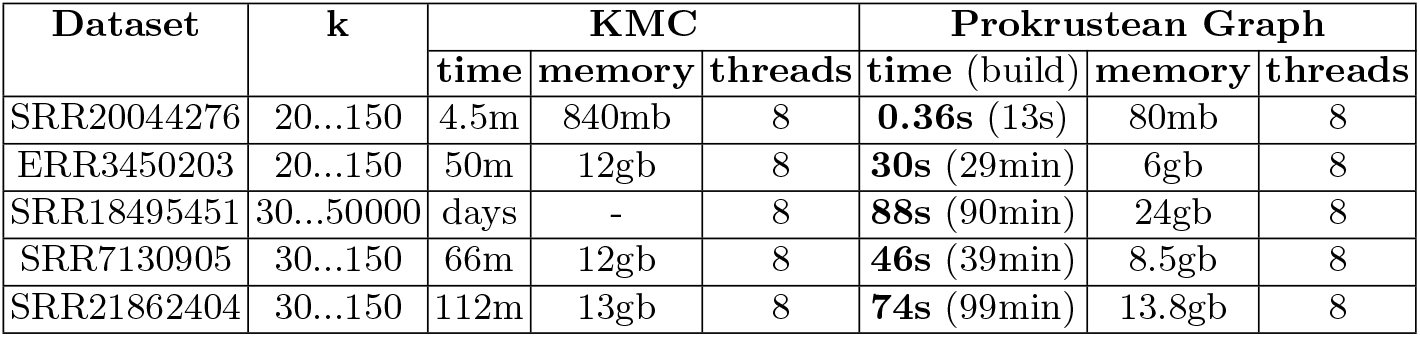
Counting distinct *k*-mers with Pro*k*rustean graphs with Algorithm S1. For comparisons, KMC had to be executed iteratively causing the computational time to increase steadily as the range of *k* increases. The running time of KMC took about 1-3 minutes for each *k*-mer size.

**Table 3:** Counting maximal unitigs with GGCAT and Algorithm S3 using the Pro*k*rustean Graph. GGCAT was executed iteratively for each *k* size, with each iteration taking approximately 1-5 minutes. The efficiency gap becomes clearer when there are many possible *k*-mer sizes, as in long reads. Both methods used 8 threads.

| Dataset | k | GGCAT |  | Prokrustean Graph |  |
| --- | --- | --- | --- | --- | --- |
|  |  | time | memory | time | memory |
| SRR20044276 | 20..150 | 13m | 310mb | **0.36s** | 70mb |
| ERR3450203 | 20..150 | 98m | 1.1gb | **21s** | 6.1gb |
| SRR18495451 | 30..50000 | days | - | **127s** | 26gb |
| SRR7130905 | 30...150 | 126m | 1.4gb | **46s** | 8.5gb |
| SRR21862404 | 30...150 | 168m | 0.8gb | **76s** | 13.8gb |

**Fig. 2:**
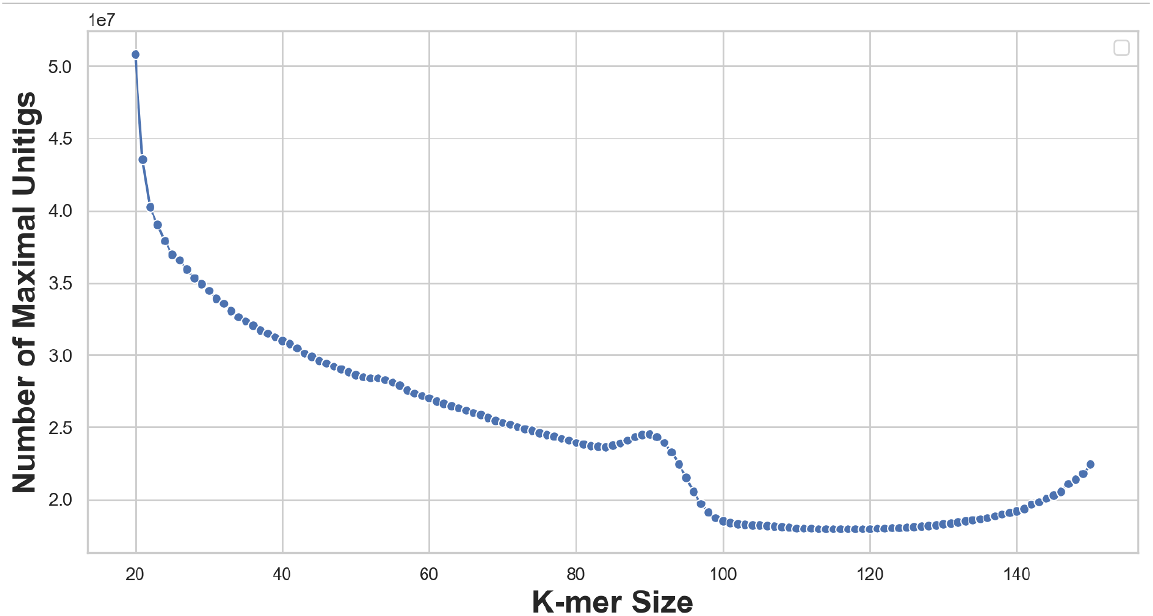
The number of maximal unitigs in de Bruijn graphs of order *k* = 20, …, 150 of metagenomic short reads (ERR3450203). A complex scenario is revealed: The decrease of the numbers fluctuates in *k* = 30, …, 60, and the peak around *k* = 90 corresponds to the sudden decrease in maximal repeats. Lastly, further disconnections increase contigs and eventually make the graph completely disconnected.

### 3.4 Computing Bray-Curtis dissimilarities for *k*_*min*_, …, *k*_*max*_

The Bray-Curtis dissimilarity is a frequency-based metric that measures the dissimilarity between biological samples. The Bray-Curtis dissimilarity for *k*-mer sets is defined as follow:

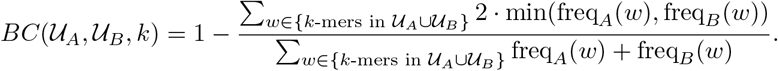

This quantity is particularly popular in metagenomics analysis [23, 36]. In each citation we found, the dissimilarity scores are presented for a single *k*-mer size. However, the values are sensitive to the choice of *k*-mer sizes, so they are often calculated with multiple *k*-mer sizes for analyses [23, 36, 47], yet most state-of-the-art tools utilize a fixed *k*-mer size [7]. Algorithm S2 computes Bray-Curtis dissimilarities for all *k*-mer sizes in *O*(|*Pairs*| *·* | 𝒢 _𝒰_ |), where 𝒢 _𝒰_ contains sequencing reads for samples, and *Pairs* is the sample pairs compared.

Figure 3 shows that dissimilarities between four example metagenome samples are not consistent in that at some *k*-mer sizes, one observed pattern of dissimilarity is completely flipped at another *k*-mer size. I.e. no *k* size is “correct”; instead, new insights are obtained from multiple *k* sizes. This task took about 10 minutes with the Pro*k*rustean graph of four samples of 12 gigabase pairs in total, and the graph construction took around 1 hour. Computing the same quantity takes around 5 to 8 minutes for a fixed *k*-mer size with Simka [7], and no library computes it across *k*-mer sizes.

**Fig. 3:**
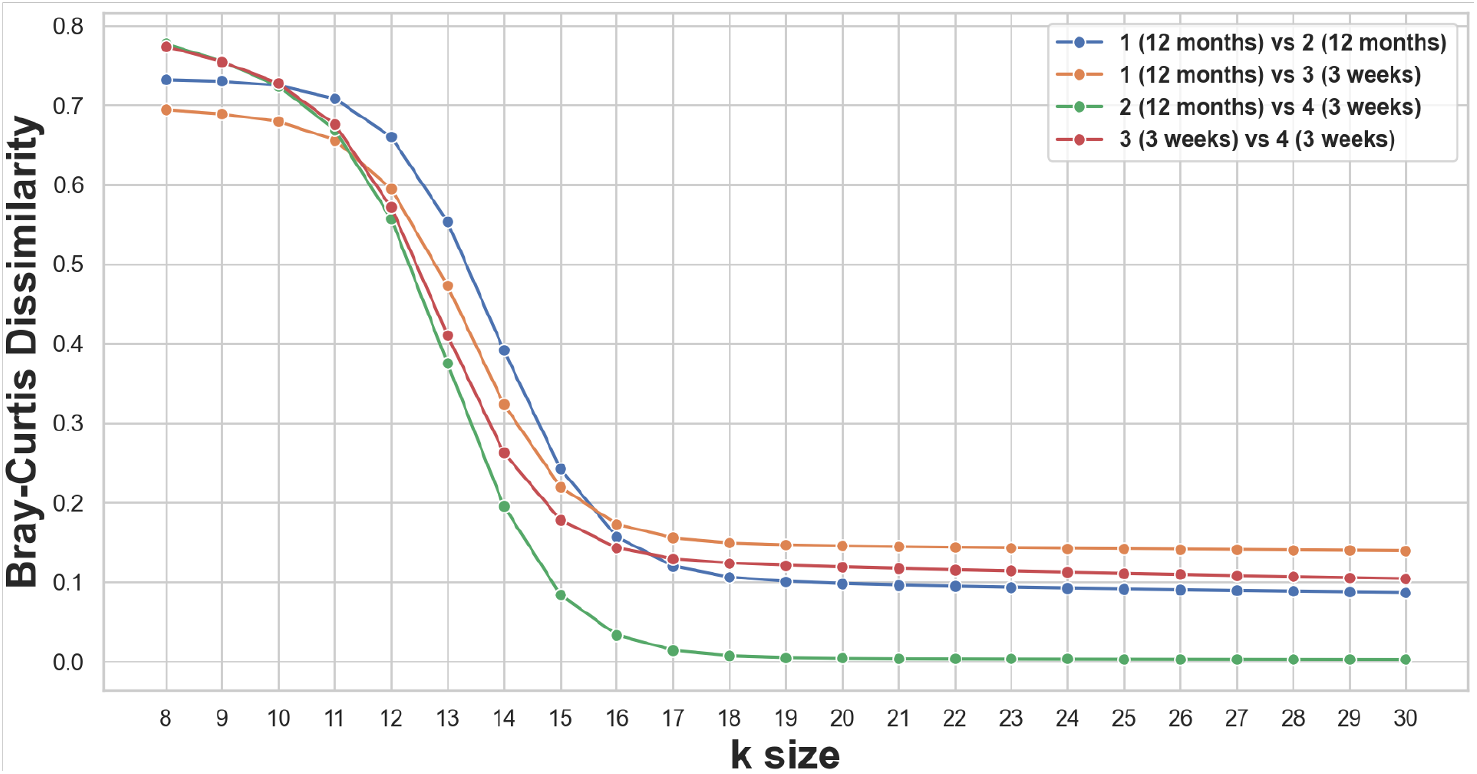
Bray-Curtis dissimilarities between four metagenomic samples computed with Algorithm S2. We used the sequencing data of four samples derived from infant fecal microbiota, which were studied in [36]. Samples include two from 12-month-old human subjects and two from 3-week-olds. The relative order between values shift as *k* changes, e.g. compared to all other dissimilarities, the dissimilarity between sample 2 and 4 is highest at *k* = 8 but lowest at *k* = 21. Note that diverse *k* sizes (10, 12, 21, 31) are actively utilized in practice.

### 3.5 Counting vertex degrees of overlap graph of threshold *k*_*min*_

Overlap graphs are extensively utilized in genome and metagenome assembly, alongside de Bruijn graphs. Although their definition appears unrelated to *k*-mers, we can view them as representing common suffix-prefix *k*-mers of sequences in 𝒰. Overlap information is particularly crucial in read classification [2, 16, 31, 41] and contig binning [32, 45] in metagenomics. These applications interpret vertex degrees as (abundant-weighted) read coverage in samples, which exhibit high variances due to species diversity and abundance variability.

A common computational challenge is the quadratic growth of overlap graphs relative to the number of reads: *O*(|𝒰 |^2^). In contrast, the Pro*k*rustean graph, which encompasses the overlap graph as a sort of hierarchical form, requires *O*(|𝒢 _𝒰_ |) space to access the structure of overlaps, enabling the efficient computation of vertex degrees.

Figure 4 is the result of computing the vertex degrees on a metagenomic short read dataset, as computed by Algorithm S4. With the Pro*k*rustean graph of 27 million short reads (4.5 gigabase pairs in total; ERR3450203), the computation used 8.5 gigabytes memory and 1 minute to count all vertex degrees.

**Fig. 4:**
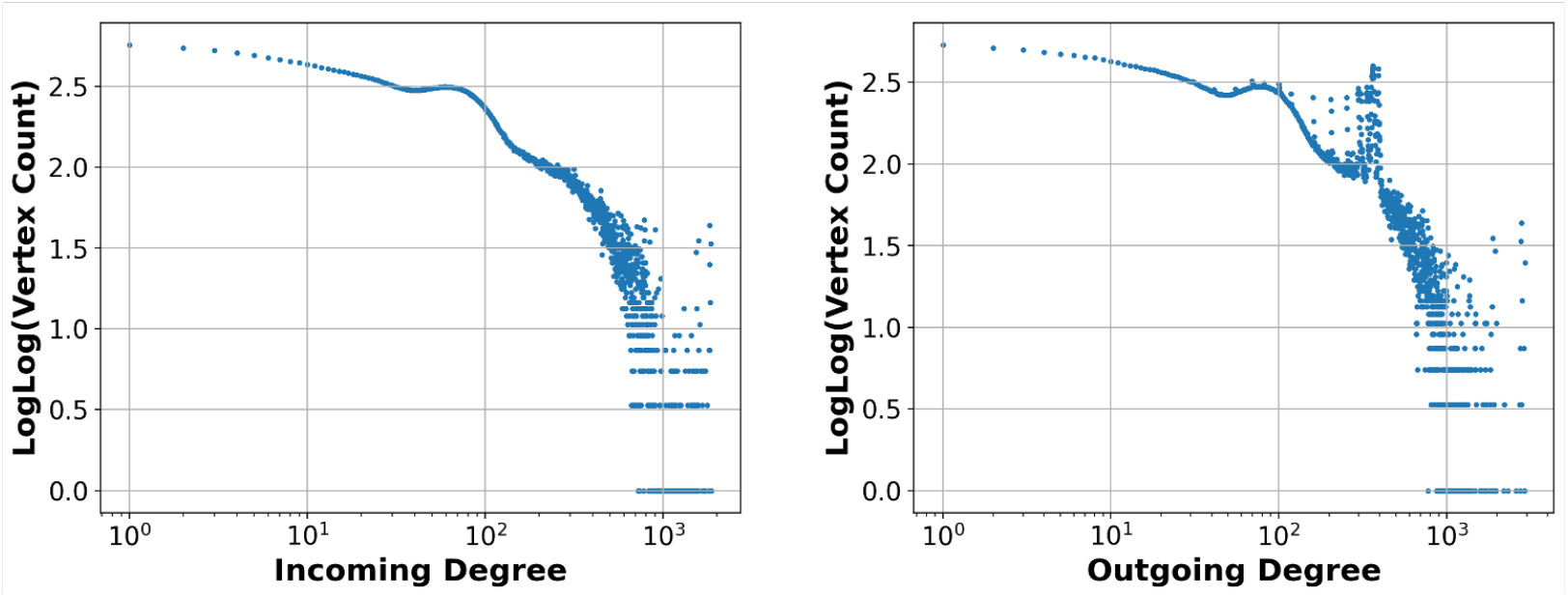
Vertex degrees of the overlap graph of metagenomic short reads (ERR3450203) computed with Algorithm S4. The number of vertices per each degree is scaled as log log *y*. There is a clear peak around 10^2^ at both incoming and outgoing degrees, which may imply dense existence of abundant species. The intermittent peaks after 10^2^ might be related to repeating regions within and across species.

## 4 Pro*k*rustean graph: A hierarchy of maximal repeats

The key idea of the Pro*k*rustean graph is to recursively capture maximal repeats by their relative frequencies of occurrences in *U*. Consider Figure 5 depicting repeats in three sequences ACCCT, GACCC, and TCCCG.

**Fig. 5:**
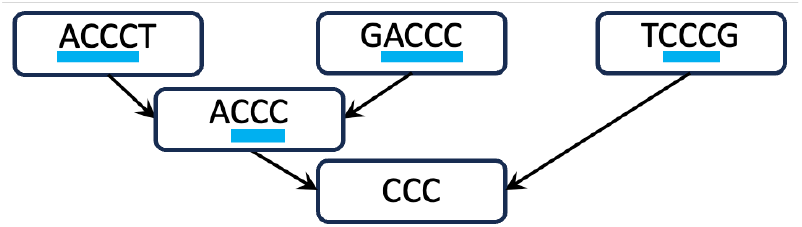
Recursive repeats. At each sequence on the top level, locally-maximal repeat regions (indicated with blue underlining) are used to denote substrings that are more frequent than its superstring. The arrows then points to a node of the substring, and the process can repeat hierarchically.

### Definition 2.

(*S, i, j*) *is <u>a locally-maximal repeat region in 𝒰</u> if:*

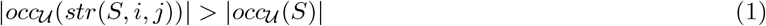

*and for each extension* (*S, i*^*′*^, *j*^*′*^) *of* (*S, i, j*),

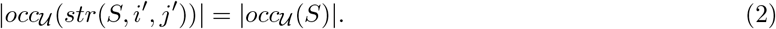

*\*\*Note that we often omit “in* 𝒰*” when the context is clear*.

A locally-maximal repeat region captures a substring that appears more frequently than the “parent” string, and any extension of the region captures a substring that appears as frequently as the parent string. Consider the previous example 𝒰 = {ACCCT, GACCC, TCCCG}. The region (ACCCT, 1, 4) is a locally-maximal repeat region capturing the substring ACCC in ACCCT. However, (ACCCT, 2, 4) is not a locally-maximal repeat region because the region capturing CCC in ACCCT can be extended to the left to capture ACCC which occurs more frequently than ACCCT.

Note, an alternative definition using substring notations, such as *locally-maximal repeats*, instead of regions, is not robust, hence the requirement for us to use region notation. Consider 𝒰= {ACCCTCCG, GCCC} and *S* :=ACCCTCCG. Observe that the region (*S*, 2, 4) capturing CCC is a locally-maximal repeat region because CCC occurs more frequently than *S* in𝒰, but its immediate left and right extensions capture ACCC and CCCT that occur the same number of times as *S*. In constrast, a subregion (*S*, 2, 3) capturing CC is not a locally-maximal repeat region because an extension (*S*, 2, 4) still captures a string (CCC) which occurs 2 times𝒰 in which is more than *S*. However, (*S*, 6, 7), which also captures CC, is a locally-maximal repeat region because |*occ*_*𝒰*_ (CC)| = 5 *>* |*occ*_*𝒰*_ (*S*)| = 1 and |*occ*_*𝒰*_ (*S*)| = |*occ*_*𝒰*_ (TCC)| = |*occ*_*𝒰*_ (CCG)|. Consequently, defining a *locally-maximal repeat* as *S*_6..7_ =CC becomes ambiguous when compared with *S*_2..3_=CC, whereas an explicit expression of a region (*S*, 6, 7) more accurately reflects the desired property of hierarchy of occurrences.

The Pro*k*rustean graph of 𝒰 is simply a graph representation of the recursive structure. Its vertexes represent the substrings and edges represent locally-maximal repeat regions. The theorem below says that all maximal repeats in 𝒰 are captured along the recursive description.

### Proposition 1 (Complete).

*Construct a string set* **R**(𝒰) *as follows:*

1. *for each locally-maximal repeat region* (*S, i, j*) *where S* ∈ 𝒰, *add str*(*S, i, j*) *to* **R**(𝒰) *and*
2. *for each locally-maximal repeat region* (*S, i, j*) *where S* ∈ **R**(𝒰), *add str*(*S, i, j*) *to* **R**(𝒰).

> *Then it follows that* **R**(𝒰) = *R* _𝒰_ *holds*.

*Proof*. Any string in **R**(𝒰) is a maximal repeat of 𝒰, so the claim is satisfied if the process captures every maximal repeat of 𝒰. Assume a maximal repeat *R* is not in **R**(𝒰). Consider any occurrence of *R* in some *S* ∈ 𝒰. Extending the maximal repeat *R* within *S* makes the number of its occurrences drop, but since *R* cannot occur as a locally-maximal repeat region by assumption, it must be included in some locally-maximal repeat region that captures a maximal repeat *R*_1_, hence *R*_1_ ∈ **R**(𝒰), and *R* is a proper substring of *R*_1_. Now consider any occurrence of *R* within *R*_1_. This argument continues recursively, capturing maximal repeats *R*_1_, *R*_2_, …, but cannot extend indefinitely as *R*_*i*+1_ is always shorter than *R*_*i*_ for every *i* ≥ 1. Eventually, *R* must be occurring as a locally-maximal repeat region of some *R*_*n*_, which is a contradiction.

Therefore, we use maximal repeats as vertices of the Pro*k*rustean graph, along with sequences, so 𝒰 ∪ ℛ _𝒰_.

### Definition 3.

*The <u>Prokrustean graph</u> of* 𝒰 *is a directed multigraph* 𝒢 _𝒰_ := (𝒱 _𝒰_, ℰ _𝒰_):

– *Vertex set* 𝒱 _*𝒰*_ : *v*_*S*_ ∈ 𝒱 _*𝒰*_ *if and only if S* ∈ 𝒰 ∪ ℛ _𝒰_.
– *Each vertex v*_*S*_ ∈ 𝒱 _*𝒰*_ *is annotated with the string size, size*(*v*_*S*_) = |*S*|.
– *Edge set* ℰ _𝒰_ : *e*_*S,i,j*_ ∈ ℰ _𝒰_ *if and only if* (*S, i, j*) *is a locally-maximal repeat region*.
– *Each edge e*_*S,i,j*_ ∈ ℰ_𝒰_ *directs from v*_*S*_ *to v*_*str*(*S,i,j*)_ *and is annotated with the interval*(*e*_*S,i,j*_) := (*i, j*), *representing the locally-maximal repeat region*.

Figure 6 visualizes how locally-maximal repeat regions are encoded in a Pro*k*rustean graph. Next, the cardinality analysis of this representation reveals promising bounds.

**Fig. 6:**
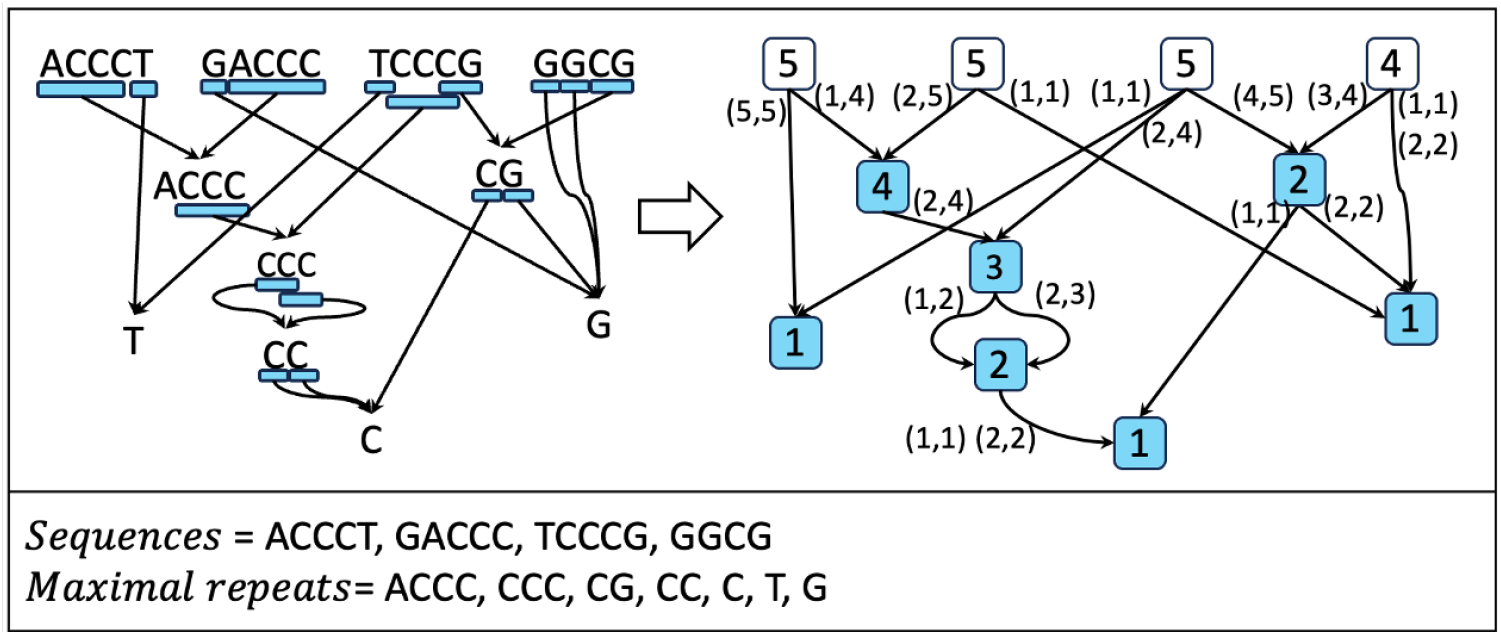
The Pro*k*rustean graph of 𝒰 ={ACCCT,GACCC,TCCCG,GGCG}. The left graph illustrates the recursion of locally-maximal repeat regions, depicted as blue regions, following the same rule described in Figure 5. The Pro*k*rustean graph on the right represents the same structure that white vertices correspond to sequences (𝒰) and blue vertices represent maximal repeats (*R*_𝒰_) found through the recursion process shown in the left graph. The strings are not stored, so integer labels store the regions and the size of the substrings.

### Theorem 2 (Compact).

|ℰ_*𝒰*_ | ≤ 2|*Σ*||𝒱_*𝒰*_ |.

*Proof*. Every vertex in *V*_*𝒰*_ has at most one incoming edge per letter extension on the right (or similarly on the left), so has at most 2|*Σ*| incoming edges. Assume the contrary: a maximal repeat appears as two different locally-maximal repeat regions (*S, i, j*) and (*S*^*′*^, *i*^*′*^, *j*^*′*^) within *S, S*^*′*^ ∈ 𝒰 ℛ ∪ 𝒰, and can extend by the same letter on the right, i.e., (*S, i, j* + 1) and (*S*^*′*^, *i*^*′*^, *j*^*′*^ + 1) capture the same string. Extending these regions further together will eventually diverge their strings, because either *S* = *S*^*′*^ or the original two regions are differently located even if *S* ≠ *S*^*′*^. Consequently, at least one of them—(*S, i, j* + 1) without loss of generality—results in a decreased occurrence count of its string as it extends. Hence, |*occ*_*𝒰*_ (*str*(*S, i, j* +1))| *>* |*occ* _𝒰_ (*S*)|. This leads to contradiction because a locally-maximal repeat region (*S, i, j*) should satisfy |*occ*_𝒰_ (*S, i, j* + 1)| = |*occ* _𝒰_ (*S*, 1, |*S*|)|. □

Assuming that |𝒰| ≪ |ℛ_𝒰_ |, as is overwhelmingly the case in practice—meaning there are significantly more maximal repeats than sequences—we derive that *O*(|𝒱_𝒰_ |) = *O*(|ℛ_𝒰_ |). Hence, *O*(|𝒢 _𝒰_ |) := *O*(|ℰ_𝒰_ |) = *O*(|*Σ*||ℛ_𝒰_ |) by the theorem. Given that |*Σ*| is typically constant in genomic sequences (eg. *Σ* = {*A, C, T, G*}), the graph’s size depends on the number of maximal repeats. Letting *N* be the accumulated sequence length of 𝒰, it is a well-known fact that |ℛ_𝒰_ | *< N* [25], so the graph grows sublinear to the input size. Furthermore, Section 3 restricted _*𝒰*_ to maximal repeats of length at least *k*_min_. Setting a practical threshold *k*_min_ allows for significantly more efficient and configurable space usage.

### 4.1 Pro*k*rustean graph construction

We leave the construction algorithm in Supplementary *Section S*3. However, we emphasize that the core insight of our study comes from the relationship between the Pro*k*rustean graph and suffix trees. It turns out that a straightforward transformation can be defined with affix trees—the union of the suffix tree of 𝒰 and the suffix tree of its reversed strings. A subset of edges of the affix tree of 𝒰 corresponds to the reversed edge set of the Pro*k*rustean graph of 𝒰. But affix trees are resource-intensive, so our final algorithm utilizes the BWT of *𝒰*, adding more intermediate steps to compensate for the non-bidirectional substring representation.

## 5 Framework: Computing *k-mer quantities* for all *k* sizes

We first describe the type of information required to efficiently compute *k-mer quantities* (defined in Theorem 1) to elucidate the uniqueness of the Pro*k*rustean graph. Specifically, why can’t other modern substring indexes be used for the computations in previous results? The answer also implies why developing efficient multi-*k k*-mer-based methods has been challenging.

### 5.1 A proxy problem: computing substring co-occurrence

There exists a theoretical void in identifying when a *k*-mer and a *k*^*′*^-mer serve similar roles within their respective substring sets. Although close *k*-mer sizes generally yield comparable outputs, dissecting this phenomenon at the level of local substring scopes is unexpectedly challenging. For instance, de Bruijn graphs constructed from sequencing reads with *k* = 30 and *k*^*′*^ = 31 display distinct yet highly similar topologies [40]. However, anyone formally articulating the topological similarity would find it quite elusive, as vertex mapping and other techniques establishing correspondences between the two graphs fail to consistently explain it.

A common underlying difficulty is that the entire *k*-mer set undergoes complete transformations as the *k*-mer size changes. Since the role of a *k*-mer is assigned within its *k*-mer set, the role of a *k*^*′*^-mer within *k*^*′*^-mers cannot be “locally” derived. To address this issue, we propose substring co-occurrence as a generalized framework for consistently grouping *k*-mers of similar roles across varying *k*-mer sizes.

#### Definition 4.

*A string S <u>co-occurs within</u> a string S*^*′*^ *in U if:*

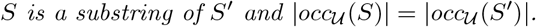

Modern substring indexes built on longest common prefixes of suffix arrays are adept at capturing co-occurrence while extending a substring. In suffix trees, a substring that extends from *S* to *S*^*′*^ along an edge—without passing through a node—indicates preservation of co-occurrence. Similarly, the BWT and its variants represent both *S* and *S*^*′*^ with the same set of suffix array intervals to imply co-occurrence. However, there has been no recognized necessity for capturing co-occurrence for substrings rather than superstrings:

#### Definition 5.

*co-substr*_*𝒰*_ (*S*) *is the set of substrings that co-occur within S*.

Modern substring indexes would fail to “smoothly” move over the order (*k*) unless *co*-*substr*_*𝒰*_ (*S*) is efficiently accessible. For instance, the variable-order de Bruijn graph and its variants extend a compact de Bruijn graph representation built on the Burrows-Wheeler Transform [12]. To address the large substring space, these solutions require non-trivial operation times for both forward moves and order changes [5, 11].

We argue the usefulness of *co*-*substr*_*𝒰*_ (*S*) with a simple yet foundational property.

#### Theorem 3 (Principle).

*For any substring S such that occ*_*𝒰*_ (*S*) *>* 0,

> *S co-occurs within exactly one sequence or maximal repeat in* 𝒰 ∪ ℛ _𝒰_.

*Proof. Assume the theorem does not hold, i*.*e*., *S co-occurs within either no string or multiple strings in U* ∪*R*_*𝒰*_. *Consider the former case: S co-occurs within no string in* 𝒰 ∪ *R*_𝒰_, *meaning S occurs more than any of its superstrings in* 𝒰 ∪ *ℛ* _𝒰_. *Given that S occurs in some string S*^*′*^∈ 𝒰 *(since occ* _𝒰_ (*S*) *>* 0*), S must be occurring more frequently than S*^*′*^ *by assumption. Then, there exists a maximal repeat R such that S is a substring of R and R is a substring of S*^*′*^, *because extending S within S*^*′*^ *decreases its occurrence at some point. Again, whenever S is a substring of a maximal repeat R, S must be occurring more frequently than R by assumption. Following the similar argument above, there exists another maximal repeat R*^*′*^ *such that S is a substring of R*^*′*^ *and R*^*′*^ *is a proper substring of R. This recursive argument will eventually terminate as R*^*′*^ *is strictly shorter than R. Thus, S co-occurs within the last maximal repeat, which contradicts the assumption.*

*Now consider the latter case where S co-occurs within at least two strings in* 𝒰 ∪ ℛ _𝒰_. *If S is extended within these two strings, either to the right or left by one step at each time, the extensions eventually diverge into two different strings since the two superstrings differ, thereby reducing the number of occurrences. Hence, S must be occurring more frequently than at least one of those two superstrings, leading to a contradiction*. ℛ _𝒰_ □

This theorem provides two insights. First, the sequences and maximal repeats 𝒰 ∪ ℛ _𝒰_ comprehensively and disjointly cover the entire substring, i.e., every substring in 𝒰 is in 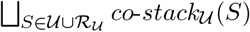. Second, by computing *co*-*substr*_𝒰_ (*S*) for every *S* ∈ 𝒰 ∪ ℛ_𝒰_, we can categorize *k*-mers of similar roles across various lengths. Therefore, we aim to design a succinct representation of *co*-*substr*_𝒰_ (*S*) that can be accessed by the Pro*k*rustean graph.

### 5.2 An algorithmic idea: co-occurrence stacks

We apply the Pro*k*rustean graph to compute *k*-mer quantities for all *k* sizes within the 𝒪 (|𝒢_𝒰_ |) time and space. Previously, a toy example in Figure 5 depicted blue regions that cover locally-maximal repeat regions. Then, given a *k*-mer size, a *complementary* region is a maximal region that covers *k*-mers that are not included in a blue region. See the red regions underlining in Figure 7. An opportunistic property is that *k*-mers in red regions co-occur within their parent strings. The following notation captures substrings in red regions.

**Fig. 7:**
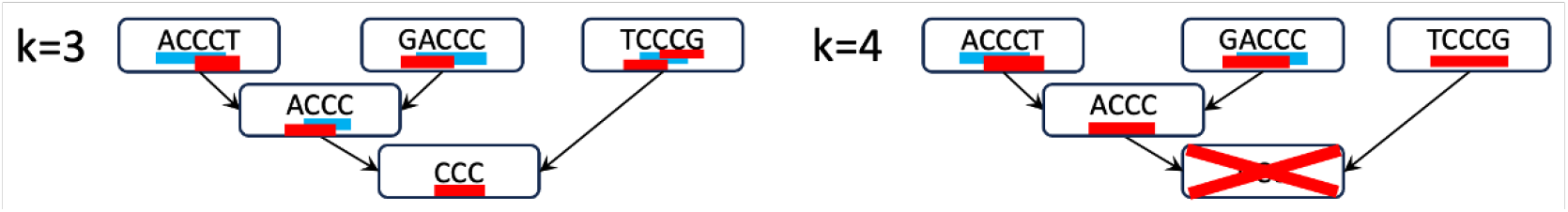
Complementary regions. Blue underlining depicts locally maximal regions, and red underlining depicts regions that complement *k*-mers not covered in blue regions, given *k* values of 3 and 4. Every *k*-mer appears exactly once within a red region. For example, in the left image, the 3-mer ACC is included in a red region within ACCCC and does not appear again in any other red region. Note that red regions can include multiple co-occurring *k*-mers. So, in the right image, both 4-mers TCCC and CCCG co-occur within TCCCG.

#### Definition 6.

*A string S <u>k-co-occurs within</u> a string S*^*′*^ *in* 𝒰 *if:*

> *every k-mer in S co-occurs within S*^*′*^ *in* 𝒰.

Furthermore, *S* <u>maximally</u> *k*-co-occurs within *S*^*′*^ in 𝒰 if no superstring of *S k*-co-occurs within *S*^*′*^ in 𝒰. The red regions in Figure 7 identify maximal *k*-co-occurring substrings of *S*^*′*^, so the Pro*k*rustean graph can be used to efficiently compute co-occurring *k*-mers of a fixed *k*-mer size. Now, we extend red regions for all possible *k* values, as shown in Figure 8. Observe that stack-like structures are formed around locally-maximal repeat regions of the string. The shape motivated the following definition:

**Fig. 8:**
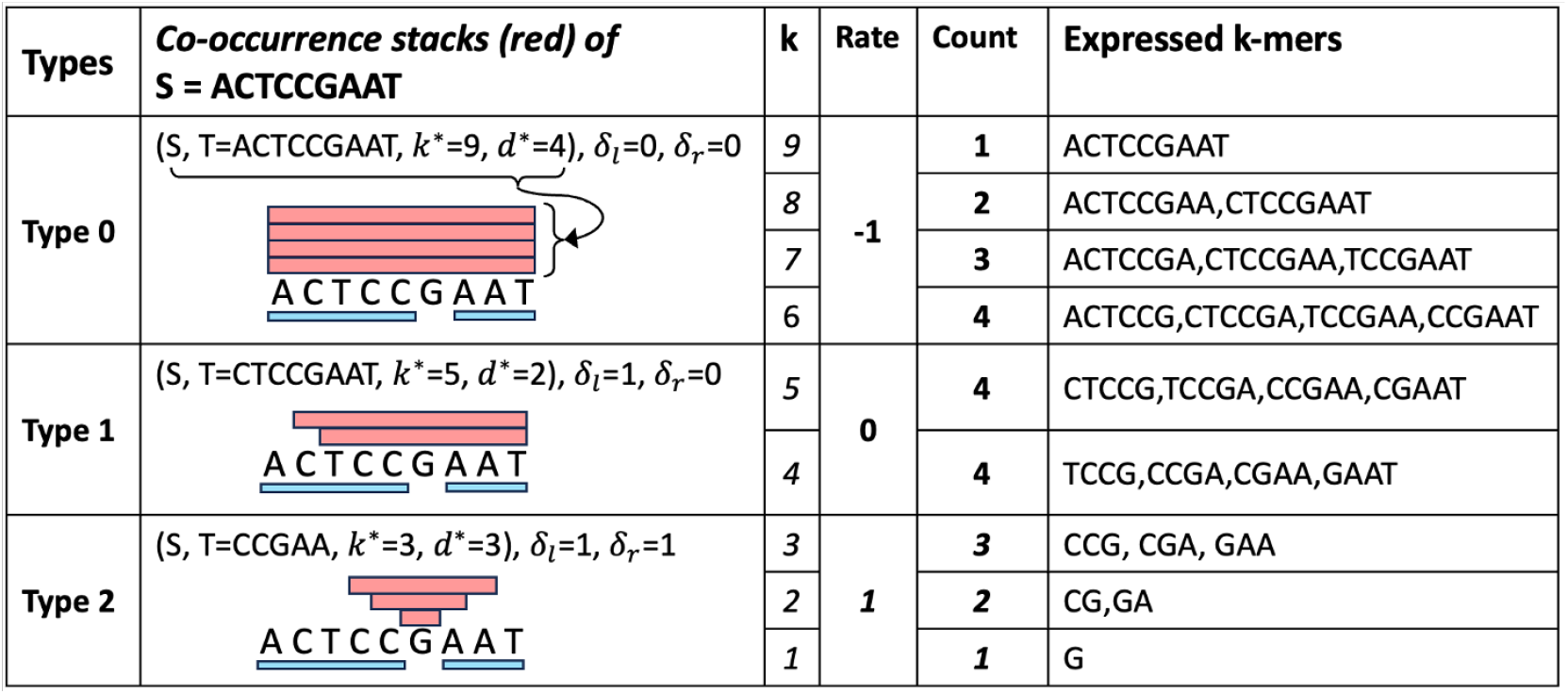
Three types of maximal co-occurrence stacks. Maximal co-occurrence stacks of *S* (sets of red rectangles) are introduced at each row for each type, given two locally-maximal repeat regions in *S* (blue rectangles). The Rate column describes the change in the count of expressed *k*-mers as the *k*-mer size increases by 1. Hence, the rate is *δ*_*l*_ + *δ*_*r*_ − 1. Also, *δ*_*l*_ and *δ*_*r*_ control the vertical shapes of stacks. For example, the Type 0 stack forms like a rectangle with *δ*_*l*_ = *δ*_*r*_ = 0, but the Type 1 stack forms like stairs on its left with *δ*_*l*_ = 1.

#### Definition 7.

*Consider two strings S and T where T is a substring of S, along with two numbers: a depth d*^∗^ *and a k-mer size k*^∗^, *each at least 1. A <u>co-occurrence stack</u> of S in U is defined as a 4-tuple:*

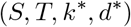

*if for each i* = 1, 2, …, *d*^∗^, *for k*_*i*_ = *k*^∗^ − (*i* − 1), *and for <u>change rates</u> δ*_*l*_ := ***1***_*T is NOT a prefix of S*_ *and δ*_*r*_ := ***1***_*T is NOT a suffix of S*_, *the following holds:*

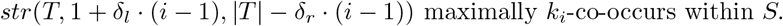

A co-occurrence stack <u>expresses</u> a *k*-mer if the *k*-mer is a substring of some *k*-co-occurring substring identified by the stack. A co-occurrence stack is <u>maximal</u> if its expressed *k*-mers are not a subset of those expressed by any other co-occurrence stack.

Let the set of all maximal co-occurrence stacks of *S* in 𝒰 as *co*-*stack*_𝒰_ (*S*). It is straightforward to check that no two stacks in *co*-*stack*_𝒰_ (*S*) express the same *k*-mer together. So, we can nicely cover *co*-*stack*_𝒰_ (*S*) with *co*-*stack*_𝒰_ (*S*). We extend the scheme to the substring space of 𝒰. Define 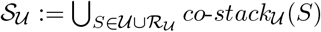.

#### Theorem 4 (Framework).

𝒮_𝒰_ *is a complete substring representation of* 𝒰 *such that every substring (any k-mer of any k value) in* 𝒰 *is expressed by exactly one stack in* 𝒮_𝒰_. |𝒮 _𝒰_ | *is bounded by O*(|𝒢_𝒰_ |), *and can be enumerated in O*(|𝒢_𝒰_ |) *time given* 𝒢_𝒰_.

*Proof. Supplementary Section S2 derives this result*. □

### 5.3 Algorithms

𝒮_𝒰_ is used in the four algorithms computing the results in Section 3. For each stack in *S*_𝒰_, constant-time operations are defined, and since |𝒮 _𝒰_ | is *O*(|𝒢_𝒰_ |), the time complexity of the algorithms is independent of the *k*-mer size interval [*k*_*min*_, *k*_*max*_].

For example, consider using the stack in the first row of Figure 8 for counting distinct *k*-mers. The co-occurrence stack (*S, T* = *ACTCCGAAT, k*^∗^ = 9, *d*^∗^ = 4) compactly represents the information of co-occurring substrings. From *k* = 6 to *k* = 9, the counts of expressed *k*-mers of *S* consistently decrease by 1, starting from 4. Therefore, for a count vector defined for the interval [*k*_*min*_, *k*_*max*_], an algorithm can implement the contribution of the stack by “adding 4 to the count vector at *k* = 6” and “recording a change rate decrease of 1 at *k* = 6 and restoring it at *k* = 9.” Repeating this process for all stacks in 𝒮_𝒰_ counts all *k*-mers in 𝒰 and takes *O*(|𝒮_𝒰_ |) time. Refer to Supplementary Algorithm S1, Algorithm S2, Algorithm S3, and Algorithm S4 for the whole contexts.

## 6 Conclusion

We have introduced the problem of computing co-occurring substrings *co*-*substr*_𝒰_ (*S*) as a proxy to analyze the influence of *k*-mer sizes in *k*-mer-based methods. The Pro*k*rustean graph facilitates access to *co*-*substr*_𝒰_ (*S*) by iterating over *co*-*stack*_𝒰_ (*S*), which is used to implement algorithms that compute *k*-mer-based quantities across a range of *k* sizes. Once the Pro*k*rustean graph is constructed using the BWT in a space-efficient manner, the computation of *k*-mer-based quantities for all *k*-mer sizes becomes roughly equivalent to scanning the graph. Furthermore, we can control the performance by choosing *k*_*min*_, the smallest maximal repeat size, so that the graph size is adjusted based on the needs of analyzed methods.

The Pro*k*rustean graph and the affix tree have intuitive relationships, as shown in Supplementary Figure S10. The contrast lies in accessing the co-occurrence structure. Suffix trees represent co-occurrence by extending substrings, functioning as a “bottom-up” approach. The Pro*k*rustean graph, however, follows a “top-down” approach, as co-occurring substrings (not superstrings) are identified in constant time. The issue with the bottom-up approach is that not all co-occurrence stacks *co*-*stack*_𝒰_ (*S*) are easily identified, even if bidirectional substring indexes are employed. For example, the Type-2 stack in Figure 8 requires information about two locally-maximal repeat regions within close proximity in a string, which is difficult to identify with substring extending operations. We therefore conjecture that exploring the space of *k*-mers with modern LCP-based substring indices presents an intrinsic challenge.

The results in Section 3.3 on counting maximal unitigs imply that a Pro*k*rustean graph can smoothly explore de Bruijn graphs of all orders. Thus, the Pro*k*rustean graph can advance the so-called variable-order scheme [11] by representing the union of de Bruijn graphs of all orders. A straightforward approach is to use the maximal co-occurrence stacks 𝒮_𝒰_ as a vertex set. We have confirmed that an edge set of size *O*(|𝒢_𝒰_ |) can represent all extensions between the stacks. Since the Pro*k*rustean graph does not grow faster than the number of maximal repeats, this new representation can serve as a compact multi-*k* assembly graph, more precisely identifying sequencing read errors, SNPs in population genomes, and genome contigs.

Thus, the open problem 5 in [40], which calls for a practical representation of variable-order de Bruijn graphs that generates assembly comparable to that of overlap graphs, can be answered with the Pro*k*rustean graph. Recall that Section 3.5 analyzed the overlap graph of 𝒰. The two popular objects used in genome assembly can be accessed together through the Pro*k*rustean graph.

There is abundant literature analyzing the influence of *k*-mer sizes in various bioinformatics tasks, but little effort has been made to derive rigorous formulations or quantities to explain these phenomena. We expect our framework to contribute as an initial step toward understanding the influence, at least at the base level of *k*-mer-based objects of sequencing data or reference genomes.

## Supplemental Material

### S1 Datasets

– Datasets mainly used in applications. SRR20044276, ERR3450203, SRR18495451, SRR21862404, SRR7130905
– Human pangenome references (Chromosome 1). AP023461.1, CH003448.1, CH003496.1, CM000462.1, CM001609.2, CM003683.2, CM009447.1, CM009872.1, CM010808.1, CM021568.2, CM034951.1, CM035659.1, CM039011.1, CM045155.1, CM073952.1, CM074009.1, CP139523.1, NC_000001.11, NC_060925.1
– Metagenome e.coli references. 3682 E. coli assemblies in NCBI circa 2020. (https://zenodo.org/records/6577997)
– Metagenome sample for Bray-Curtis, referring to [36]. ERS11976829, ERS11976830, ERS11976565, ERS11976566

### S2 Full framework

We complete the framework in Section 5. Refer to the definitions and figure below which were already introduced in the main text.

#### Definition S8.

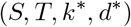

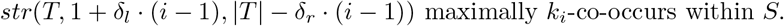

A co-occurrence stack <u>expresses</u> a *k*-mer if the *k*-mer is a substring of some *k*-co-occurring substring identified by the stack. A co-occurrence stack is <u>maximal</u> if its expressed *k*-mers are not a subset of those expressed by any other co-occurrence stack. It is clear that any *k*-mer expressed in a co-occurrence stack of *S* co-occurs within *S*. So, maximal co-occurrence stacks of *S* collectively express *co*-*substr*_𝒰_ (*S*).

#### Definition S9.

*co-stack*_𝒰_ *(S) is the set of all maximal co-occurrence stacks of S in* 𝒰.

A maximal co-occurrence stack of *S* is <u>type-0</u> if the substring *T* is *S* itself, <u>type-1</u> if *T* is a proper prefix or suffix of *S*, and <u>type-2</u> otherwise. The numeral in each type’s name indicates the number of locally-maximal repeat regions intersecting the regions of substrings identified by the co-occurrence stack. Refer to Figure 8 for intuition on the type names. We further dig into the computational part regarding the types.

**Fig. S9:**
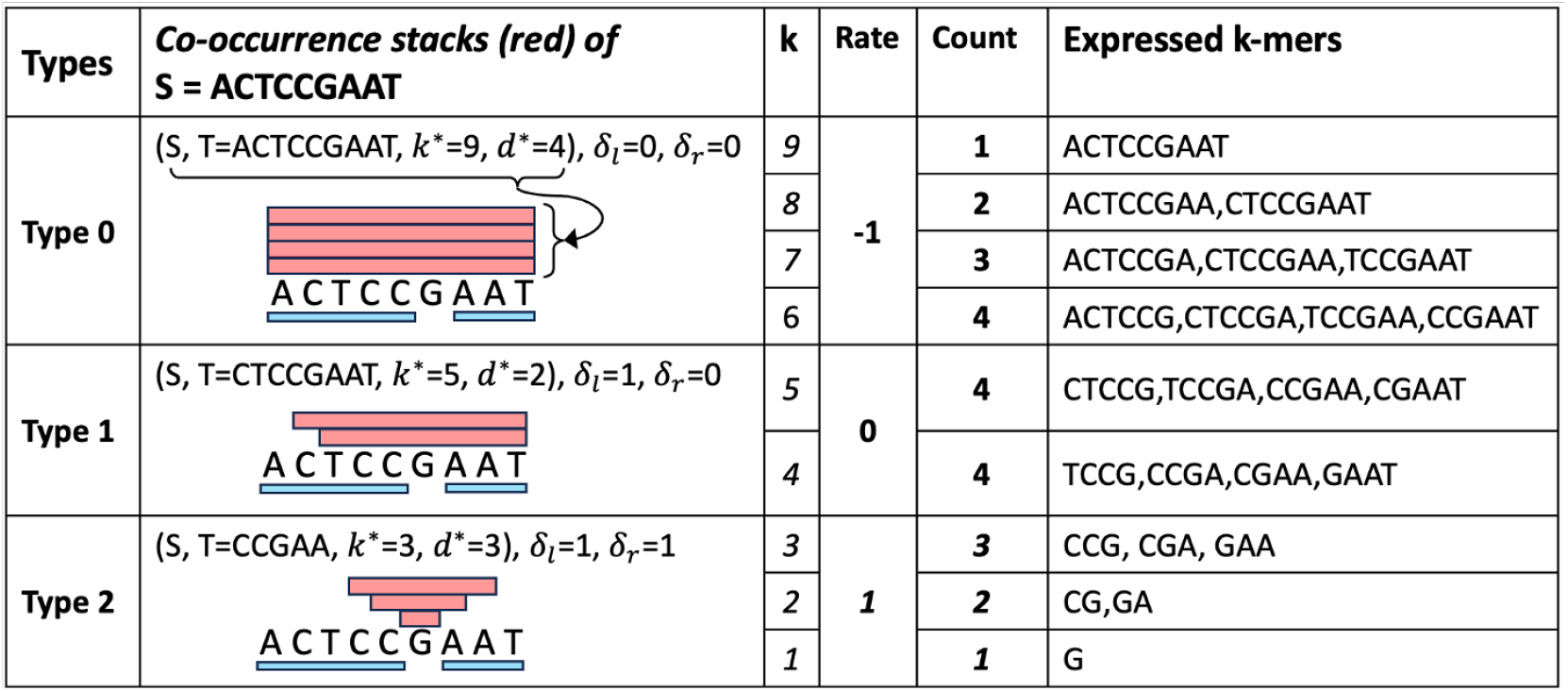
Three types of maximal co-occurrence stacks.

#### Proposition 2.

*A string S maximally k-co-occurs within a string S*^*′*^ *in U if and only if its region in S*^*′*^ *intersect locally-maximal repeat regions by less than k* − 1, *and on both sides, either extends to an end of S*^*′*^ *or intersects a locally-maximal repeat region by exactly k* − 1.

*Proof. The forward direction is straightforward: if S k-co-occurs within S*^*′*^, *its region cannot intersect any locally-maximal repeat region by k at any case, and if the intersection is under k* − 1 *on its edge, by extending S and making the intersection k* − 1, *another superstring of S k-co-occurs within S*^*′*^.

*For the reverse direction, assume a region satisfies the conditions. A k-mer appearing in the region cannot occur more than S, as it would mean the region intersects a locally-maximal repeat region by at least k, so S*^*′*^ *k-co-occurs within S. S*^*′*^ *is also maximal because extending its region increases the size of the intersection to more than k* − 1. □

#### Proposition 3.

*A maximal k-co-occurring substring T and a maximal k*^*′*^*-co-occurring substring T* ^*′*^ *of S are identified by the same maximal co-occurrence stack if and only if they share identical boundary conditions: on both sides, they either extend to the end of S or intersect the same locally-maximal repeat region by k* − 1 *and k*^*′*^ − 1, *respectively*.

*Proof. Consider maximal k-co-occurring substring T and k*^*′*^*-co-occurring substring T* ^*′*^ *of S where k < k*^*′*^. *The forward direction is straightforward; a maximal co-occurrence stack* (*S, S*^*′*^, *k*^∗^, *d*^∗^), *which identifies T and T, identifies S*^*′*^ *as a k*^∗^*-co-occurring substring of S by definition, and S*^*′*^ *either extends to an end of S or intersects an locally-maximal repeat region by k*^∗^ − 1 *on both sides according to Proposition 2. Then, as T and T* ^*′*^ *are identified as the* (*k*^∗^ −*k* + 1)*-th and* (*k*^∗^ −*k*^*′*^ + 1)*-th elements by the stack, respectively, the rates δ*_*l*_ *and δ*_*r*_ *derived from the stack ensure T and T* ^*′*^ *sharing the equivalent side conditions with S*^*′*^ *as described in Proposition 2*.

*For the reverse direction, consider three types of co-occurrence stacks: 1. T* = *T* ^*′*^ = *S, i*.*e*., *they extend to both ends of S. Then a type-0 stack identifies them. Whenever S k-co-occurs within S, the same holds for k* + 1 *and above until* |*S* |. *Therefore*, (*S, T* := *S, k*^∗^ := |*S*|, *d*^∗^) *with some maximal d*^∗^ *identifies T and T* ^*′*^.

*2. T and T* ^*′*^ *are proper prefixes of S. Then a type-1 stack identifies them. The condition says there is a locally-maximal repeat region* (*S, i, j*) *intersecting the regions of T and T* ^*′*^ *in S by k* − 1 *and k*^*′*^ − 1, *respectively. Without loss of generality, let the intersection be on the left of* (*S, i, j*). *Then, a k*^∗^*-co-occurring prefix of S with k*^∗^ := | (*S, i, j*) | *intersects* (*S, i, j*) *by k*^∗^− 1; *if not, there must be a k*^∗^*-mer occurring more than* |*occ*_𝒰_ (*S*)| *starting between positions 1 and i* − 1. *But k-mers of T start in the positions too, so T does not k-co-occur within S, which leads to contradiction. Thus*, (*S, T* := *str*(*S*, 1, *j*− 1), *k*^∗^ := | (*S, i, j*) |, *d*^∗^) *is a type-1 stack that identifies both T and T* ^*′*^.

*3. T and T* ^*′*^ *are neither prefixes nor suffixes. Then a type-2 stack identifies them. There are two locally-maximal repeat regions* (*S, i, j*) *and* (*S, i*^*′*^, *j*^*′*^) *that intersect the regions of T and T* ^*′*^ *by k*− 1 *and k*^*′*^− 1, *respectively. Assuming* (*S, i, j*) *is smaller, a similar argument as in the previous paragraph shows that a k*^∗^*-co-occurring substring intersects both* (*S, i, j*) *and* (*S, i*^*′*^, *j*^*′*^) *by k*^∗^ − 1 *given k*^∗^ := |(*S, i, j*)|, *so* (*S, T* := *str*(*S, j* − *k*^∗^ + 2, *i*^*′*^ + *k*^∗^ − 2), *k*^∗^ := |(*S, i, j*)|, *d*^∗^) *identifies T and T* ^*′*^. □

This proposition implies that either zero, one, or two locally-maximal repeat regions in *S* characterize a co-occurrence stack by identifying substrings of identical side conditions. This corresponds to the classifications—type-0, type-1, and type-2. The proposition also implies that the number of maximal co-occurrence stacks of *S* does not grow faster than the number of locally-maximal repeat regions in *S*.

#### Theorem S5.

*co-stack*_*𝒰*_ (*S*) *completely and disjointly cover co-occurring substrings of S, i*.*e*.,

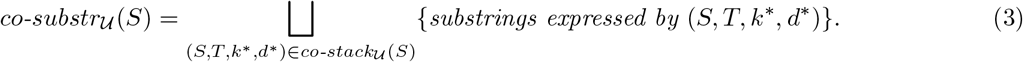

*Also*,

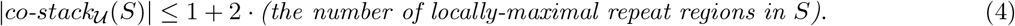

*Hence, given* 𝒢 _𝒰_, *enumerating co-stack*_*𝒰*_ (*S*) *takes O*(|*outgoing edges of v*_*S*_ *in* ℰ _𝒰_ |) *time*.

*Proof. co*-*stack*_*𝒰*_ (*S*) completely and disjointly covers *co*-*substr* _𝒰_ (*S*): any *k*-mer occurring in 𝒰 co-occurs within an exactly one string *S* in 𝒰 ∪ ℛ _𝒰_ by Theorem 3. The *k*-mer extends to a unique superstring that maximally *k*-co-occurs within *S* because appearing in two distinct maximally *k*-co-occurring substrings imply multi-occurrence of the *k*-mer in *S*. The superstring is identified by a unique maximal co-occurrence stack by Proposition 3. So, the *k*-mer is expressed by exactly one maximal co-occurrence stack.

To demonstrate the inequality, first, consider a type-0 stack on a string *S*, where all identified substrings are *S* itself by definition, i.e., extend to ends of *S*. A single type-0 stack identifies them all by Proposition 3. Hence, just one type-0 stack identifies them all by Proposition 3.

Next, for a type-1 or type-2 maximal co-occurrence stack (*S, T, k*^∗^, *d*^∗^), we show its longest identified substring always intersects some locally-maximal repeat region of length *k*^∗^ by exactly *k*^∗^− 1. Since a proper type-1 stack identifies every *k*-co-occurring prefix (or suffix) of *S* whose region intersects the same locally-maximal repeat region by *k*− 1, by Proposition 3, the largest order *k*^∗^ works too. Then, *k*^∗^ should be the length of the locally-maximal repeat region; otherwise, we can find a (*k*^∗^ + 1)-co-occurring prefix (or suffix) of *S*, denoted *T* ^*′*^, intersecting the locally-maximal repeat region by *k*^∗^, so (*S, T* ^*′*^, *k*^∗^ + 1, *d*^∗^ + 1) is a proper co-occurrence stack, resulting in the original stack non-maximal. Similarly, a type-2 stack identifies substrings that intersect two locally-maximal repeat regions on both sides, so checking the length of the smaller region be *k*^∗^ permits a similar deduction.

Note that each left and right side of a locally-maximal repeat region of length *l* intersect the region of at most one maximally *l*-co-occurring substring by exactly *l*− 1. Since such maximal intersection happens with every type-1 or type-2 maximal co-occurrence stack at least once, the number of such stacks is bounded by twice the number of locally-maximal repeat regions, thus is 2(the number of outgoing edges of *v*_*S*_ in ℰ_*𝒰*_) in total.

Lastly, enumerating *co*-*stack*_*𝒰*_ (*S*) is efficient. A type-0 stack is uniquely defined for a string *S* where the depth *d*^∗^ is one more than the length of the largest locally-maximal repeat region. Each side of a locally-maximal repeat region maximally intersects the largest substring identified in at most one stack, either type-1 or type-2. Without loss of generality, a type-1 stack is defined on the left side of a locally-maximal repeat region if it is the largest one among those found on its left, while a type-2 stack pairs with the nearest locally-maximal repeat region of at least the same size on its left. These properties can be verified by a single scan of the locally-maximal repeat regions of a string. □

Lastly, this scheme is extended to the entire substring space.

#### Proposition 4.

*The number of all maximal co-occurrence stacks in U is O*(|*𝒢*_*𝒰*_)|. *Precisely*,

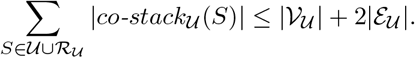

*Proof*. This is implied by the inequality Equation (4). The term |𝒱_𝒰_ | accounts for the constant contribution of 1 from each *S* ∈ 𝒰 ∪ ℛ_𝒰_, and 2|ℰ_𝒰_ | corresponds to the right term because ℰ_𝒰_ represents the total set of locally-maximal repeat regions in 𝒰 ∪ ℛ_𝒰_. □

Since *k*-mers expressed by each stack of *S* inherit characteristics of *S*, such as frequency and extensions within𝒰, algorithms can leverage this to compute *k-mer quantities*. We introduce four applications motivated by biological problems that benefit from this approach.

#### S2.1 Application: Counting distinct *k*-mers of all *k* sizes

The core idea leverages the change rate of *k*-mers within each co-occurrence stack, thereby eliminating the need to consider *k*-mer counts individually. The number of *k*-mers expressed by each type-0, 1, or 2 co-occurrence stack changes by -1, 0, or +1 as *k* increases by 1, respectively, as detailed in the Rate column of Figure S9. This property is utilized in lines 4–7 in Algorithm S1. Recalling that |*co*-*stack*_*𝒰*_ (*S*)| is bounded by 1 + 2 *×* (the number of outgoing edges of *v*_*S*_) in Theorem S5, the nested loop in lines 2-3 runs in *O*(|*𝒢*_*𝒰*_ |) time.

##### Algorithm S1 Count *k*-mers of a range of *k* sizes

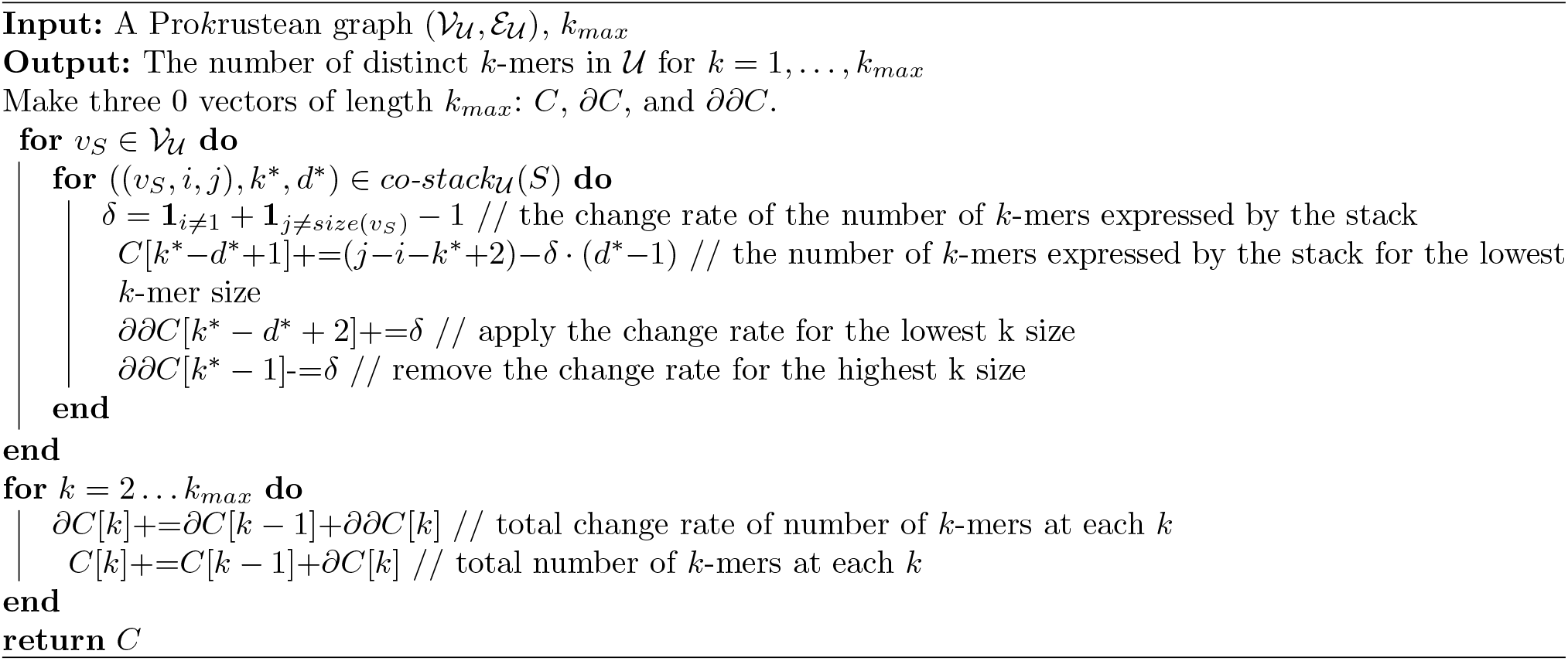

#### S2.2 Application: Computing Bray-Curtis dissimilarities of all *k* sizes

Recall the definition of the Bray-Curtis dissimilarity for *k*-mer sets.

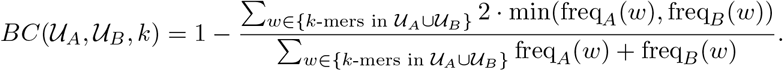

We can compute the Bray-Curtis dissimilarity for all *k*-mer sizes of samples A and B using 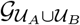, i.e., the Pro*k*rustean graph constructed from the union of their sequence sets *𝒰*_*A*_ and *𝒰*_*B*_. For each 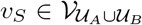 such that *S* ∈ *𝒰*_*A*_ ∪ *𝒰*_*B*_, whether *S* is from *A* or *B* can be recorded while constructing 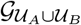. Then, freq_*A*_(*w*) is 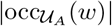, accessed through some *v*_*S*_ ∈ 𝒱_*𝒰*_ satisfying *w* co-occurs within *S*. The algorithm is contained in Algorithm S2.

##### Algorithm S2 Compute Bray-Curtis dissimilarities of a range of *k* sizes

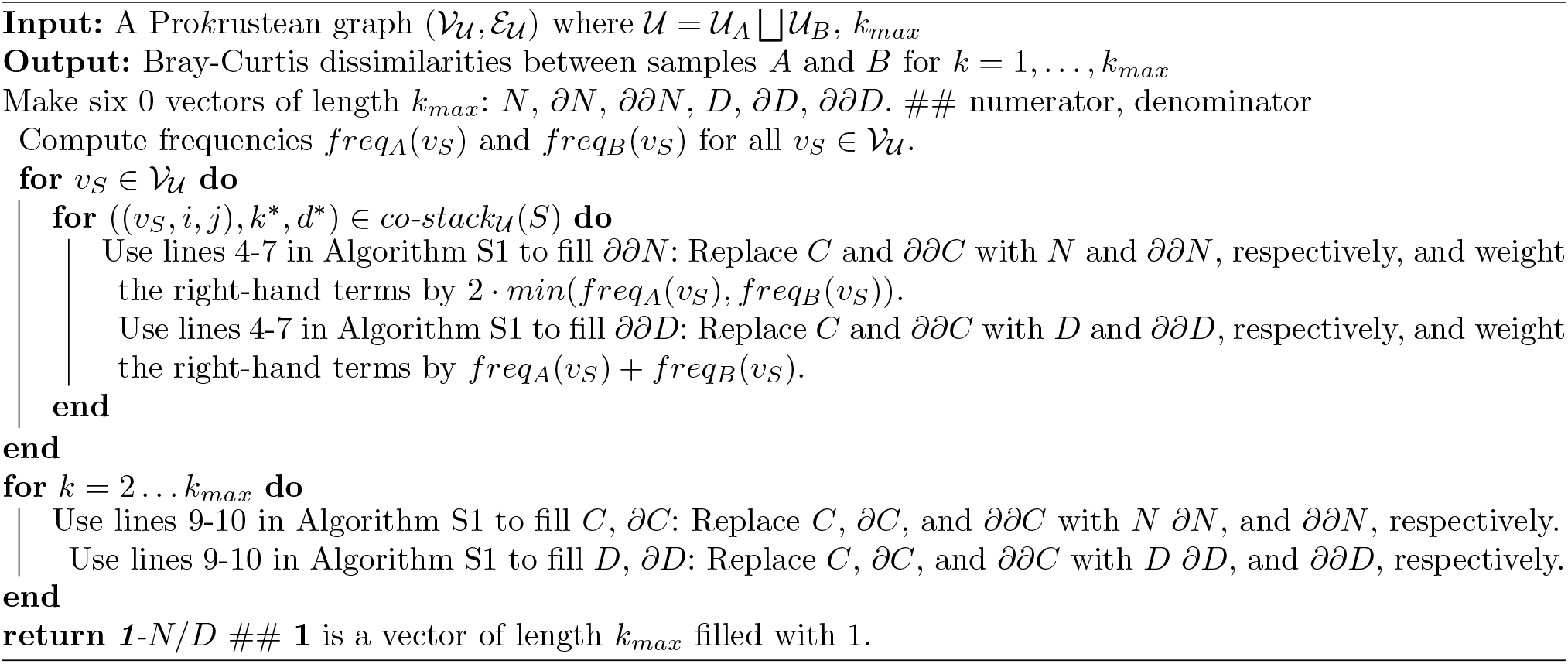

Recall that the value assignments in line 5 and 6 utilized co-occurrence stacks for counting *k*-mers in Algorithm S1. The values *min*(*freq*_*A*_(*v*_*S*_), *freq*_*B*_(*v*_*S*_)) and *freq*_*A*_(*v*_*S*_) + *freq*_*B*_(*v*_*S*_) are assigned instead of a constant 1, because they grow linearly to the number of co-occurring *k*-mers weighted by the frequency-based values. Consequently, vectors *N* and *D* contains values of the numerator and denominator part of the Bray-Curtis dissimilarities for all *k*-mer sizes.

Typically, biological experiments compute these values for pairs of multiple samples. The proposed algorithm can be extended to consider multiple samples, requiring *O*(|samples| ^2^ *·*|𝒢 _𝒰_ |) time for 𝒰 the union of all sample read sets.

#### S2.3 Application: Counting maximal unitigs of all *k* sizes

Recall that a maximal unitig is a simple path in the de Bruijn graph that cannot be extended further. The idea for computing the number of maximal unitigs is that they are implied by tips, convergences, and divergences, which are indicated by type-0 or 1 co-occurrence stacks. For instance, consider a string *S* in 𝒰∪ ℛ_𝒰_ with multiple right extensions, such as C and T. If a suffix region in *S* of length *l* is an locally-maximal repeat region, it suggests that a *k*-co-occurring suffix of *S* where *k > l* corresponds to a vertex in the respective de Bruijn graph of order *k* with right extensions C and T. Therefore, identifying the locally-maximal repeat regions on suffixes and prefixes of strings in 𝒰∪ ℛ_𝒰_ is sufficient. This idea is summarized in Algorithm S3.

##### Algorithm S3 Count maximal unitigs of de Bruijn graphs for a range of *k* sizes

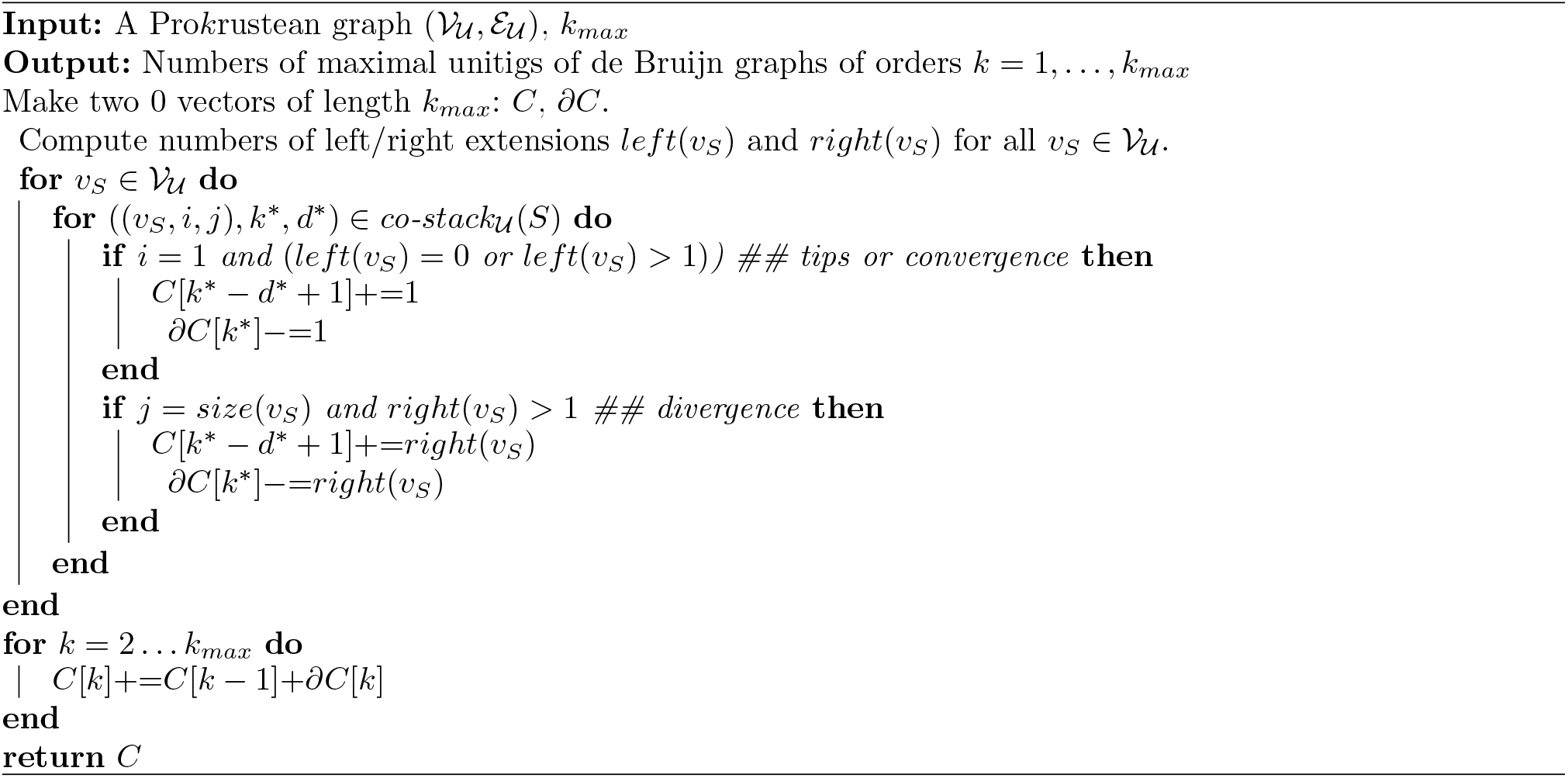

This algorithm runs in *O*(|𝒢 _𝒰_|) time, but can be accelerated in practice. Storing left(*v*_*S*_) and right(*v*_*S*_) in the Pro*k*rustean graph is not burdensome due to the small alphabet size (|*Σ*|), which allows them to be represented with 1 byte. Also, co-occurrence stacks should not be enumerated at line 4. Instead, it is enough to find two *k*^∗^ values that are exactly one larger than locally-maximal repeat regions each located at the leftmost and rightmost positions in *S*, respectively. Then, *C*[*k*^∗^ − *d*^∗^ + 1]+=1 and ∂*C*[*size*(*v*_*S*_)]−=1 computed for those two *k*^∗^ values are straightforwardly equivalent to running line 4 to 10. So, assuming left(*v*_*S*_) and right(*v*_*S*_) can be accessed in constant time, Algorithm S3 can run in *O*(|*𝒱*_𝒰_ |) time. So, the topological characteristics of de Bruijn graphs of all orders can be analyzed within time linear to the number of maximal repeats.

#### S2.4 Application: Computing vertex degrees of overlap graph

*V U*

The overlap graph of 𝒰 has the vertex set 𝒰, and an edge (*v*_*S*_, *v*_*S*_*′*) if and only if a suffix of *S* matches a prefix of *S*^*′*^. Define *pref* (*v*_*R*_ ∈ 𝒱_𝒰_) with a substring *R*, as the number of *R* occurring as a prefix in strings in, which is easily computed through the recursive structure.

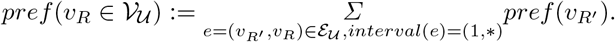

Once *pref* is computed, the outgoing degree of a vertex *v*_*S*_ is the sum of all *pref* (*v*_*R*_) such that *R* is a prefix maximal repeat of *S*. This computation requires two recursions of depth-first explorations of the Pro*k*rustean graph, resulting in an overall time complexity of *O*(|*𝒢*_*𝒰*_ |). Refer to Algorithm S4 for a brief description.

##### Algorithm S4 Count vertex outgoing degrees of the overlap graph

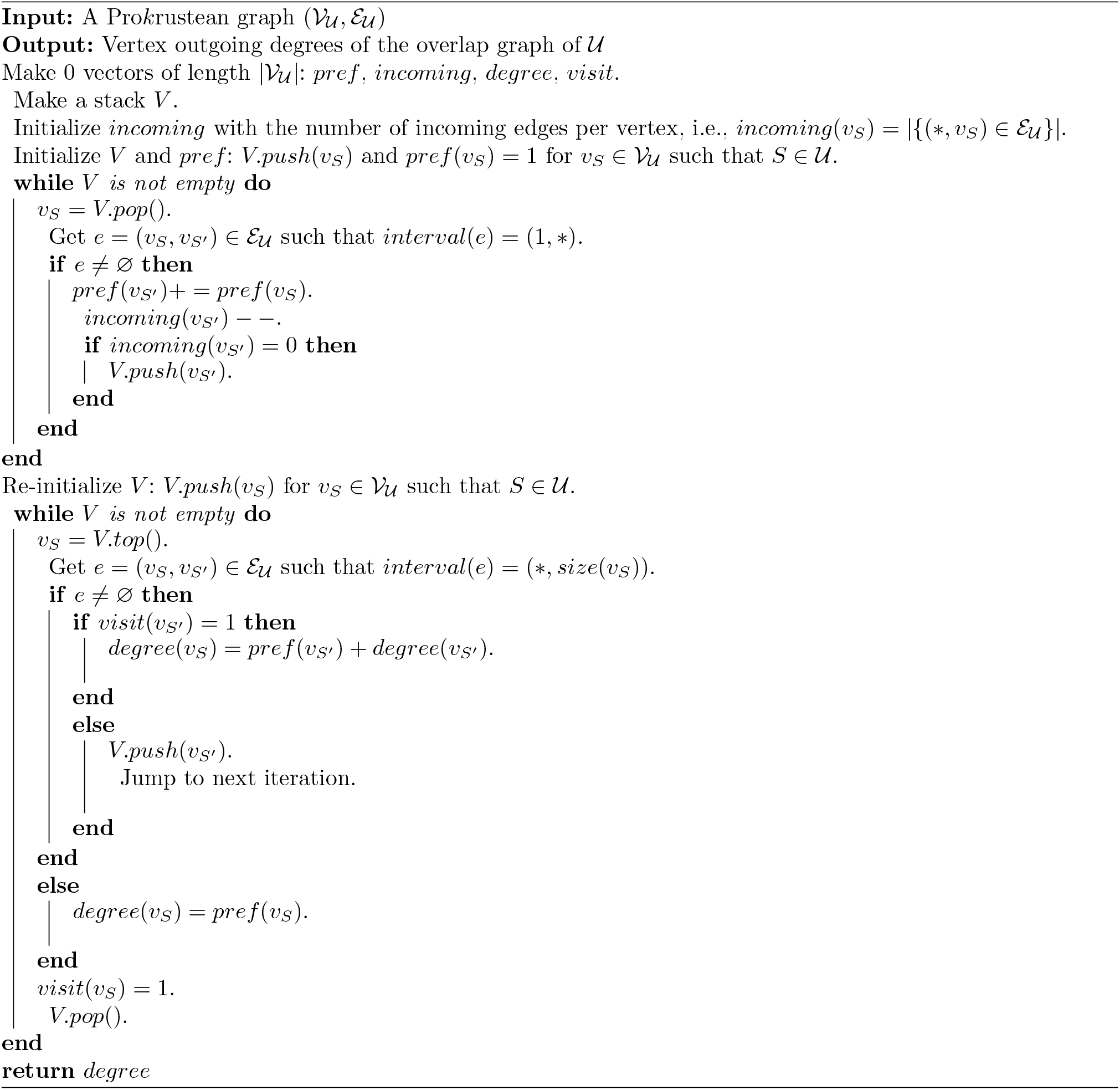

### S3 Pro*k*rustean graph construction

In this section, we describe the Pro*k*rustean construction processes, starting with a simpler to describe, but less efficient approach before proceeding to the full implementation.

Recall that Section 5.1 addressed the limitations of LCP-based substring indexes in computing *co*-*substr*_*𝒰*_ (*S*), which motivated the adoption of the Pro*k*rustean graph. LCP-based representations inherently provide a “bottom-up” approach and are at best able to access incoming edges of *v*_*S*_ in the Pro*k*rustean graph. In contrast, the Pro*k*rustean graph supports a “top-down” approach by accessing *co*-*substr*_𝒰_ (*S*) through co-occurrence stacks whose number depends on the outgoing edges of *v*_*S*_.

We start with a more straightforward, but less efficient approach: Section S3.1 employs affix trees—the union of the suffix tree of 𝒰 and the suffix tree of its reversed strings—to extract the Pro*k*rustean graph. Although affix trees allow straightforward operations supporting bidirectional extensions above suffix trees, they are resource-intensive in practice. Section S3.2 refines the approach using the BWT of𝒰, incorporating an intermediate model to compensate the bidirectional context lost in moving away from affix trees. Section S3.3 then briefly introduce the implementation, covering recent advancements in BWT-based computations supporting it.

#### S3.1 Extracting Pro*k*rustean graph from two suffix trees

For ease of explanation, we use two suffix trees to represent the affix tree. Let 𝒯 _𝒰_ denote the generalized suffix tree constructed from𝒰. The reversed string of a string *S* is denoted by *rev*(*S*), and the reversed string set *rev*(𝒰) is {*rev*(*S*) | *S* ∈ 𝒰}. A node in a tree is represented as *node*(*S*) if the path from the root to the node spells out the string *S*, with the considered tree being always clear from the context. We assume affix links are constructed, which map between *node*(*S*) in 𝒯 _𝒰_ and *node*(*rev*(*S*)) in 𝒯 _*rev*(𝒰)_ if both nodes exist in their respective trees.

Additionally, a string preceded/followed by a letter represents an extension of the string with that letter, as in *σS* or *Sσ*. Similarly, concatenation of two strings *S* and *S*^*′*^ can be written as *SS*^*′*^. An asterisk (*) on a substring denotes any number (including zero) of letters extending the substring, as in *S*∗.

**Vertex set** Collecting maximal repeats of 𝒰 requires information about |*occ*_*𝒰*_ (*σR*) | and |*occ*_*𝒰*_ (*Rσ*) | for each *σ* ∈ *Σ*, for any considered substring *R*. Most substring indexes built above LCPs provide access to occurrences through the LCP intervals, and the number of occurrences is represented by the length of the corresponding LCP intervals. The reverse representation 𝒯 _*rev*(𝒰)_ is not required yet, implying that the same information is accessible via single-directional indexes like the BWT in the next section.

**Edge set** A more nuanced approach is required to extract the edge set of a Pro*k*rustean graph. Since an edge in a suffix tree basically implies co-occurrence, it sometimes directly indicates a locally-maximal repeat region, providing a subset of substrings that co-occur within their extensions. That is, an edge from *node*(*S*) to *node*(*S*^*′*^) in 𝒯 _𝒰_ implies that *S* may represent a locally-maximal repeat region in *S*^*′*^ capturing a prefix *S* of *S*^*′*^. However, different scenarios may arise—sometimes the reverse direction should be considered, or both directions, or neither. These variations depend on the combinations of letter extensions around the substring. We present a theorem that delineates these cases, visually explained in Figure S10 for intuition.

We define terms distinguishing occurrence patterns of extensions. A string *R* is <u>left-maximal</u> if |*occ*_𝒰_ (*σR*)| *<* |*occ*_𝒰_| (*R*) for every *σ* ∈ *Σ*, and <u>left-non-maximal with</u>*<u>σ</u>* if |*occ*_𝒰_ (*σR*) | = |*occ*_𝒰_ (*R*) |, and <u>right-maximality</u> and right-non-maximality are symmetrically defined.

##### Theorem S6.

*Consider a maximal repeat R* ∈ ℛ _𝒰_ *and letters σ*_*l*_, *σ*_*r*_ ∈ *Σ. The following three cases define a bijective correspondence between selected edges in the suffix trees* (𝒯 _𝒰_ *and* 𝒯_*rev*(𝒰)_) *and the edges* (ℰ _𝒰_) *of the Prokrustean graph of* 𝒰.

***Case S6*.*1*** *(Prefix Condition) The following three statements are equivalent*.

a. *Rσ*_*r*_ *is left-maximal*.
b. *There exists a string T of form Rσ*_*r*_∗ *in* 𝒰 ∪ ℛ_𝒰_ *such that an edge from node*(*R*) *to node*(*T*) *is in* 𝒯 _𝒰_.
c. *There exists an edge e*_*T,i,j*_ *in* ℰ_𝒰_ *satisfying str*(*T, i, j* + 1) = *Rσ*_*r*_ *and i* = 1.

***Case S6*.*2*** *(Suffix Condition) The following three statements are equivalent*.

a. *σ*_*l*_*R is right-maximal*.
b. *There exists a string T of form* ∗*σ*_*l*_*R in* 𝒰 ∪ ℛ_𝒰_ *such that an edge from node*(*rev*(*R*)) *to node*(*rev*(*T*)) *is in T*_*rev*(*𝒰*)_.
c. *There exists an edge e*_*T,i,j*_ ∈ ℰ_*𝒰*_ *satisfying str*(*T, i* − 1, *j*) = *σ*_*l*_*R and j* = |*T* |.

**Fig. S10:**
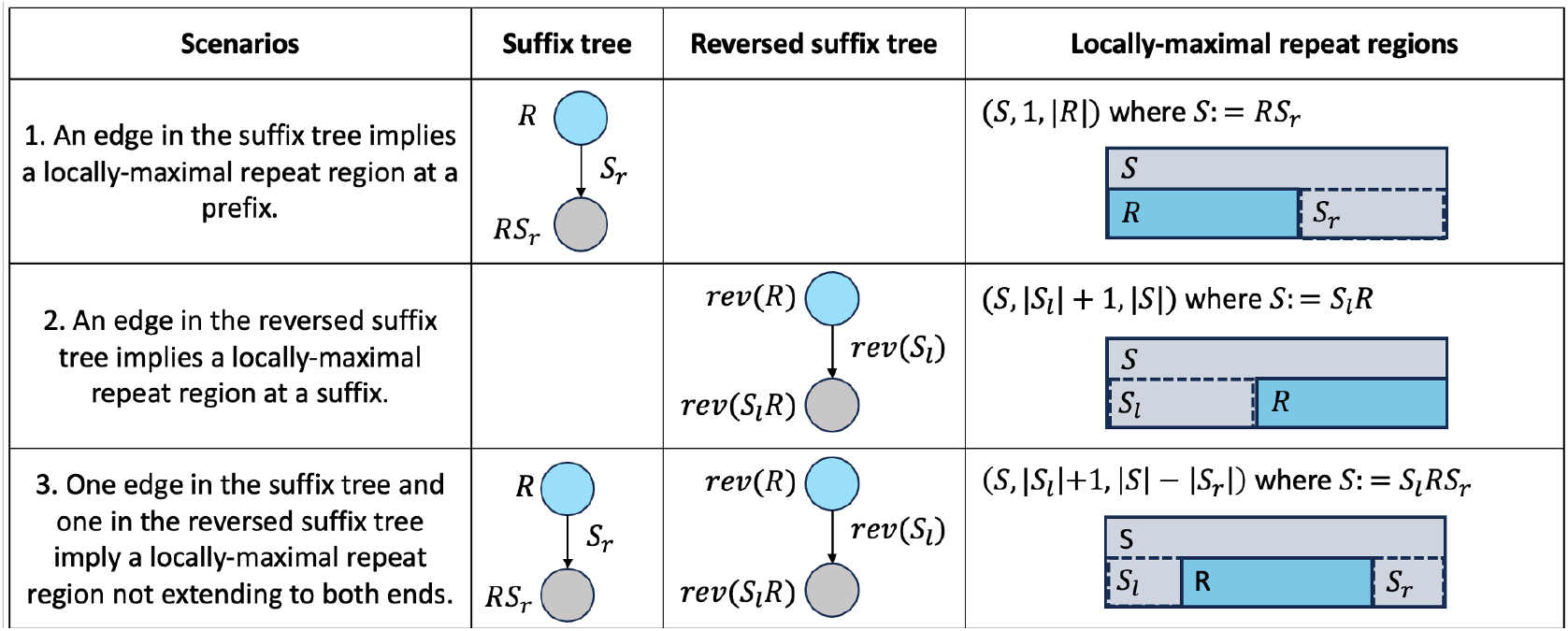
Deriving locally-maximal repeat regions from edges of suffix trees. From a node of a maximal repeat *R* or *rev*(*R*), the extension *S* is identified by adding strings on edge annotations. Then, the three scenarios depict how each edge(s) locates each locally-maximal repeat region that captures *R* within *S*.

***Case S6*.*3*** *(Interior Condition) The following three statements are equivalent*.

a. *Rσ*_*r*_ *is left-non-maximal with σ*_*l*_ *and σ*_*l*_*R is right-non-maximal with σ*_*r*_.
b. *There exist strings S*_*l*_ *and S*_*r*_, *and T* := *S*_*l*_*RS*_*r*_ *of form* ∗*σ*_*l*_*Rσ*_*r*_∗ *in* 𝒰 ∪ ℛ_𝒰_ *such that an edge from node*(*R*) *to node*(*RS*_*r*_) *is in* 𝒯 _𝒰_ *and an edge from node*(*rev*(*R*)) *to node*(*rev*(*S*_*l*_*R*)) *is in* 𝒯*rev*(𝒰).
c. *There exists an edge e*_*T,i,j*_ *in* ℰ_𝒰_ *satisfying str*(*T, i* − 1, *j* + 1) = *σ*_*l*_*Rσ*_*r*_.

*Proof*. Case S6.1 (*a*) =⇒ (*b*): We proceed arguing in the contrapositive. Assume *T* := *RS*_*r*_ ∉ 𝒰 ∪ ℛ _𝒰_ which is non-maximal on either left or right. Observe that the definition of the construction of the suffix tree 𝒯 _𝒰_ implies |*occ*_*𝒰*_ (*Rσ*_*r*_)| = |*occ*_*𝒰*_ (*RS*_*r*_)| and *RS*_*r*_ being right-maximal. Since *RS*_*r*_ is right-maximal, *RS*_*r*_ must be left-non-maximal with some *σ* ∈ *Σ*, so |*occ*_*𝒰*_ (*σRS*_*r*_)| = |*occ*_*𝒰*_ (*RS*_*r*_)|. Since |*occ*_*𝒰*_ (*Rσ*_*r*_)| = |*occ*_*𝒰*_ (*RS*_*r*_)|, |*occ*_*𝒰*_ (*σRS*_*r*_)| = |*occ*_*𝒰*_ (*RS*_*r*_)| = |*occ*_*𝒰*_ (*Rσ*_*r*_)| meaning *Rσ*_*r*_ is left-non-maximal with some *σ*.

Case S6.1 (*b*) =⇒ (*c*): Since *R, T* ∈ 𝒰 ∪ ℛ_𝒰_ holds and *T* [|*R*| + 1] = *σ*_*r*_, showing (*T*, 1, |*R*|) is a locally-maximal repeat region is enough to show *e*_*T*,1,|*R*|_ satisfies the statement. Let *T* := *RS*_*r*_. First, the region’s string *R* occurs more than *RS*_*r*_ because *R* is a maximal repeat. Next, as (*RS*_*r*_, 1, |*R*|) is a prefix region, every extension includes the smallest extension (*RS*_*r*_, 1, |*R*| + 1), so the string of any of its extensions occurs at least |*occ*_𝒰_ (*Rσ*_*r*_)|, but 𝒯 _𝒰_ implies |*occ*_𝒰_ (*Rσ*_*r*_)| = |*occ*_𝒰_ (*RS*_*r*_)|, so the extensions co-occur within *RS*_*r*_. Hence, (*T*, 1, |*R*|) is a locally-maximal repeat region and *str*(*T*, 1, |*R*| + 1) = *Rσ*_*r*_.

Case S6.1 (*c*) =⇒ (*a*): Since *T*, 1, |*R*| is a locally-maximal repeat region, its extension captures *Rσ*_*r*_, so |*occ*_𝒰_ (*Rσ*_*r*_)| = |*occ*_𝒰_ (*T*)| holds. Also, *T* ∈ 𝒰 ∪ ℛ_𝒰_ implies *T* is left- and right-maximal, so |*occ*_𝒰_ (*σT*)| *<* |*occ*_𝒰_ (*T*)| holds for any *σ* ∈ *Σ*. Therefore, |*occ*_𝒰_ (*σRσ*_*r*_)| *<* |*occ*_𝒰_ (*Rσ*_*r*_)| for any *σ* ∈ *Σ*, hence *Rσ*_*r*_ is left-maximal.

**Case S6.2 is symmetrically argued in line with Case S6.1.

Case S6.3 (*a*) =⇒ (*b*): The non-maximalities in (*a*) imply |*occ*_𝒰_ (*σ*_*l*_*R*)| = |*occ*_𝒰_ (*Rσ*_*r*_)| = |*occ*_𝒰_ (*σ*_*l*_*Rσ*_*r*_)|. Also, by co-occurrences implied by two suffix trees, |*occ*_𝒰_ (*S*_*l*_*R*)| = |*occ*_𝒰_ (*σ*_*l*_*R*)| and |*occ*_*𝒰*_ (*RS*_*r*_)| = |*occ*_𝒰_ (*Rσ*_*r*_)| hold. Hence, |*occ*_*𝒰*_ (*S*_*l*_*R*)| = |*occ*_𝒰_ (*S*_*l*_*RS*_*R*_)| = |*occ*_𝒰_ (*RS*_*R*_)| is satisfied, meaning *S*_*l*_*RS*_*R*_ is left- and right-maximal that *T* := *S*_*l*_*RS*_*R*_ ∈ 𝒰 ∪ ℛ _𝒰_.

Case S6.3 (*b*) =⇒ (*c*): The maximal repeat *R* occurs more than *S*_*l*_*RS*_*r*_ and the string of any extension occurs at least |*occ* _𝒰_ (*σ*_*l*_*R*)| or |*occ* _𝒰_ (*Rσ*_*r*_)| times, while |*occ* _𝒰_ (*σ*_*l*_*S*)| = |*occ* _𝒰_ (*S*_*l*_*SS*_*R*_)| = |*occ* _𝒰_ (*SS*_*R*_)| is implied by the two suffix trees. Hence, (*T, i, j*) := (*S*_*l*_*RS*_*r*_, |*S*_*l*_| + 1, |*S*_*l*_| + |*R*|) is a locally-maximal repeat region such that *str*(*T, i* − 1, *j* + 1) = *σ*_*l*_*Rσ*_*r*_.

Case S6.3 (*c*) =⇒ (*a*): This statement is trivial because the locally-maximal repeat region implies |*occ* _𝒰_ (*σ*_*l*_*R*)| = |*occ*_*𝒰*_ (*S*_*l*_*RS*_*R*_)| = |*occ* _𝒰_ (*Rσ*_*r*_)|, hence |*occ* _𝒰_ (*σ*_*l*_*R*)| = |*occ*_*𝒰*_ (*Rσ*_*r*_)| = |*occ* _𝒰_ (*σ*_*l*_*Rσ*_*r*_)| holds, which implies non-maximalities on both sides of *R*.

The computation of extracting the edge set of the Pro*k*rustean graph is realized by collecting incoming edges for each vertex: Explore 𝒯 _*𝒰*_ to identify each maximal repeat *R* ∈ ℛ _𝒰_, and verify the maximality conditions in items labeled (*a*) in the theorem. Upon satisfying (*a*) of any case, use the corresponding (*b*) to identify a superstring *T* from the suffix trees 𝒯 _𝒰_ and 𝒯_*rev*(𝒰)_. The corresponding (*c*) then confirms that the region of *R* within *T* is indeed a locally-maximal repeat region in 𝒰. The precise region (*T, i, j*) is inferred from the decomposition of *T* outlined in (*b*). For instance, if *T* is *S*_*l*_*R* as in Case S6.2, then (*T, i, j*) is the suffix region (*T*, |*S*_*l*_| + 1, |*T* |), so an edge from *v*_*T*_ to *v*_*R*_ is labeled (|*S*_*l*_| + 1, |*T* |).

This approach covers every edge in the Pro*k*rustean graph. Observe that items labeled (*c*) distinguish locally-maximal repeat regions by the letters on immediate left or right. Each region is unique with respect to the letter extension, as previously established in the proof of Theorem 2. Hence, for each maximal repeat *R*, the conditions met in (*a*) have bijectively mapped incoming edges to *v*_*R*_, thus the entire edge set ℰ_𝒰_ is identified through Theorem S6.

We have leveraged a bidirectional substring index, but if only one direction is supported, i.e., 𝒯_*rev*(𝒰)_ is unavailable, finding *T* as outlined in (*b*) becomes challenging. This limitation with single-directional substring indexes motivates the development of an enhanced algorithm.

#### S3.2 Extracting Pro*k*rustean graph from Burrows-Wheeler transform

This section addresses two issues of the previous computation when implemented in practice. Firstly, suffix tree representations generally consume substantial memory, which hinders scalability when handling large genomic datasets such as pangenomes. Secondly, bidirectional indexes are more space-consuming, expensive to build, and less commonly implemented than single-directional indexes. Instead, BWTs enable traversal of the same suffix tree structure using sublinear space relative to the input sequences, and are supported by a range of actively improved construction algorithms.

Since the bidirectional extensions in items of (*b*) in Theorem S6 are not directly applicable to the BWT or other single-directional substring indexes, we utilize a subset of occurrences of maximal repeats as an intermediate representation of locally-maximal repeat regions. This approach necessitates a consistent way of utilizing suffix orders within the generalized suffix arrays of 𝒰.

Before describing the construction of the Pro*k*rustean from a BWT, we need the following definitions.

##### Definition S10.

*Let the rank of a region* (*S, i, j*) *be the rank of the suffix of S starting at i in the generalized suffix array of* 𝒰, *which is implied by a substring index being used. Then, the <u>first occurrence</u> of a string R, occ*_*𝒰*_ (*R*)_***1***_, *is the lowest ranked occurrence among occ*_*𝒰*_ (*R*).

Note that suffix ordering is consistent only within a sequence but varies across sequences depending on implementations. For example, among BWTs that employ different strategies, some consider the global lexicographical order, yielding *occ*_*𝒰*_ (*AA*)_**1**_= (CCAA, 3, 4) for 𝒰 = {GGAA, CCAA}, while others impose a strict order on input sequences, such as = {GGAA#_1_, CCAA#_2_}, resulting in *occ*_𝒰_ (*AA*)_**1**_ = (GGAA, 3, 4). This variability is extensively discussed in [18]. In any scenario, the identical Pro*k*rustean graph will be generated as long as the first occurrences are consistently considered.

The following definition of projected occurrences considers specific occurrences of maximal repeats, as an intermediate representation of the edges of the Pro*k*rustean graph.

##### Definition S11.

*ProjOcc*_*𝒰*_ *is a subset of occurrences of strings in* 𝒰 ∪ ℛ _𝒰_, *where elements are specified as follows:*

1. *For every sequence S* ∈ 𝒰, *the entire region of the sequence, i*.*e*., (*S*, 1, |*S*|) ∈ *ProjOcc*_*𝒰*_.
2. *For every maximal repeat R* ∈ ℛ 𝒰 *and letters σ*_*l*_, *σ*_*r*_ ∈ *Σ*,

1. *If Rσ*_*r*_ *is left-maximal, then* (*S, i, j* − 1) ∈ *ProjOcc*_𝒰_ *given* (*S, i, j*) := *occ*_𝒰_ (*Rσ*_*r*_)_***1***_.
2. *If σ*_*l*_*R is right-maximal, then* (*S, i* + 1, *j*) ∈ *ProjOcc*_𝒰_ *given* (*S, i, j*) := *occ*_𝒰_ (*σ*_*l*_*R*)_***1***_.
3. *If Rσ*_*r*_ *is left-non-maximal with σ*_*l*_ *and σ*_*l*_*R is right-non-maximal with σ*_*r*_, *then* (*S, i* + 1, *j* − 1) ∈ *ProjOcc*_𝒰_ *given* (*S, i, j*) := *occ*_𝒰_ (*σ*_*l*_*Rσ*_*r*_)_***1***_.

**Fig. S11:**
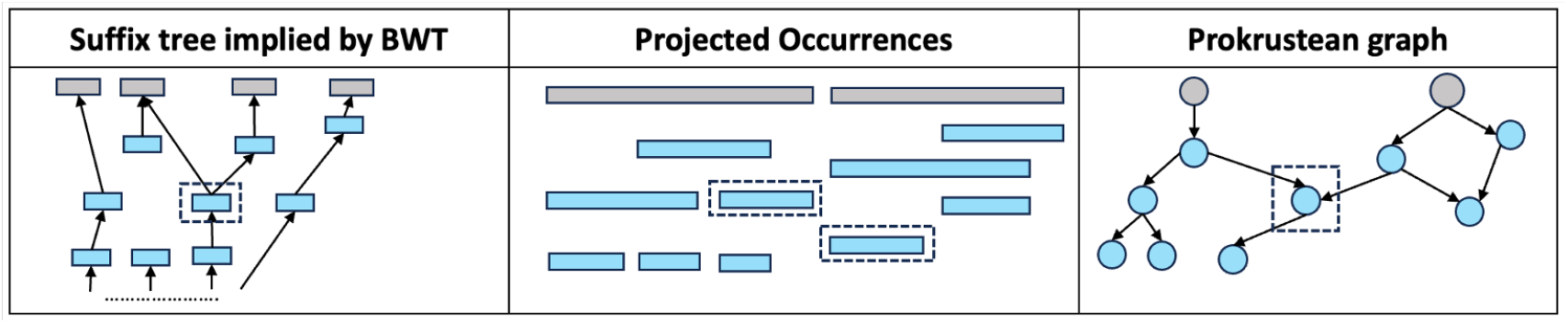
Construction of the Pro*k*rustean graph via projected occurrences. Gray vertices, depicted as either rectangles or circles, represent sequences in *𝒰*, while blue vertices denote maximal repeats of *𝒰*. The suffix tree is shown flipped to better illustrate the correspondence, providing intuition for the “bottom-up” approach. Dotted rectangles in each diagram refer to a maximal repeat *R*. First occurrences of substrings like *σ*_*l*_*R* and *Rσ*_*r*_ are identified through the construction of *ProjOcc*_𝒰_. Then, the nested structure of projected occurrences derive incoming edges of *v*_*R*_ in the Pro*k*rustean graph.

Refer to Figure S11 for intuition that *ProjOcc*_*𝒰*_ is indirectly deriving the locally-maximal repeat regions in 𝒰 ∪ ℛ _𝒰_. Although the same maximality conditions are used in both Theorem S6 and the definition of *ProjOcc*_𝒰_, the occurrences in *ProjOcc*_*𝒰*_ are organized in a nested manner, allowing locally-maximal repeat regions to be inferred through their inclusion relationships. So, *ProjOcc*_𝒰_ is a projected image of the Pro*k*rustean graph of 𝒰. The decoding rule for *ProjOcc*_𝒰_, necessary for reconstructing the Pro*k*rustean graph, is articulated in the following proposition and theorem.

The proposition below is useful to prove the main theorem.

##### Proposition 5.

*For every maximal repeat R* ∈ ℛ _𝒰_, *occ*_*𝒰*_ (*R*)_**1**_ *is in ProjOcc*_𝒰_.

*Proof*. If a maximal repeat is a whole sequence in 𝒰, it is trivially in *ProjOcc*_𝒰_. So, assume every occurrence of *R* in 𝒰 is extendable by some letters.

Let (*S, i, j*) := *occ*_*𝒰*_ (*R*)_**1**_, and define *σ*_*l*_ := *S*[*i* − 1] and/or *σ*_*r*_ := *S*[*j* + 1]. One of *σ*_*l*_ or *σ*_*r*_ might not be defined, so consider (*S, i* − 1, *j*) = *occ*_*𝒰*_ (*σ*_*l*_*R*)_**1**_ without loss of generality.

If *σ*_*l*_*R* is right-maximal, then *occ*_𝒰_ (*R*)_**1**_ trivially belongs to *ProjOcc*_𝒰_ by 2-(2) of Definition S11. Otherwise, *σ*_*l*_*R* is right-non-maximal with some *σ*_*r*_, so (*S, i, j* + 1) = *occ*_𝒰_ (*Rσ*_*r*_)_**1**_ must hold.

Consequently, whether *Rσ*_*r*_ is left-maximal or left-non-maximal with *σ*_*l*_, *occ*_𝒰_ (*R*)_**1**_ ∈ *ProjOcc* _𝒰_ holds because conditions 2-(1) or 2-(3) of Definition S11 apply, respectively. The symmetric argument assuming (*S, i, j* + 1) = *occ*_*𝒰*_ (*Rσ*_*r*_)_**1**_ confirms the same results. □

The following theorem elaborates the decoding rule. Let the relative occurrence of (*S, i, j*) within (*S, i*^*′*^, *j*^*′*^) be (*str*(*S, i*^*′*^, *j*^*′*^), *i* − *i*^*′*^ + 1, *j* − *i*^*′*^ + 1) defined only if (*S, i, j*) ⊊ (*S, i*^*′*^, *j*^*′*^).

##### Theorem S7.

*e*_*T,p,q*_ ∈ ℰ _𝒰_, *i*.*e*., (*T, p, q*) *is a locally-maximal repeat region in some T* ∈ 𝒰 ∪ ℛ_𝒰_ *if and only if there exist* (*S, i, j*), (*S, i*^*′*^, *j*^*′*^) ∈ *ProjOcc*_𝒰_ *satisfying:*

– (*S, i, j*) ⊊ (*S, i*^*′*^, *j*^*′*^), *and*
– *the relative occurrence of* (*S, i, j*) *within* (*S, i*^*′*^, *j*^*′*^) *is* (*T, p, q*), *and*
– *no region* (*S, x, y*) ∈ *ProjOcc*_𝒰_ *satisfies* (*S, i, j*) ⊊ (*S, x, y*) ⊊ (*S, i*^*′*^, *j*^*′*^).

*Proof*. The forward direction is composed of three parts. We show there exists some (*S, i, j*) ∈*ProjOcc*_𝒰_ corresponding to (*T, p, q*) that is specified by Theorem S6, and then show there exists some (*S, i*^*′*^, *j*^*′*^) ∈ *ProjOcc*_𝒰_ such that the relative occurrence of (*S, i, j*) in the region is (*T, p, q*). Lastly, the third statement is shown.

First, there exists an occurrence (*S, i, j*) ∈ *ProjOcc*_𝒰_ that captures the same string and shares some left or right extension with (*T, p, q*). (*T, p, q*) must be satisfying exactly one of three items labeled (*c*) in Case S6.1, Case S6.2, and Case S6.3 conditioned by extensions of (*T, p, q*). Then the corresponding item of (*a*) describes a maximality condition around *R* that is equally described in 2-(1), 2-(2), and 2-(3) in Definition S11 to derive elements in *ProjOcc*_*𝒰*_. So, there is always some (*S, i, j*) in *ProjOcc*_*𝒰*_ chosen for (*T, p, q*) by the agreement in letter extensions, i.e., either *S*[*i* − 1] = *T* [*p*− 1] or *S*[*j* + 1] = *T* [*q* + 1] or both holds.

Next, there exists some (*S, i*^*′*^, *j*^*′*^) such that the relative occurrence of (*S, i, j*) within (*S, i*^*′*^, *j*^*′*^) is (*T, p, q*). The construction of (*S, i, j*) showed either *S*[*i*− 1] = *T* [*p* − 1] or *S*[*j* + 1] = *T* [*q* + 1] holds, so assume *S*[*i*− 1] = *T* [*p* − 1] without loss of generality. Since (*T, p, q*) is a locally-maximal repeat region, *str*(*T, p* − 1, *q*) co-occurs within *T* and hence *str*(*S, i* − 1, *j*) co-occurs within *T*. Therefore, (*S, i, j*) can be extended to some (*S, i*^*′*^, *j*^*′*^) so that *T* is captured and *i* is at *i*^*′*^ + *p* − 1 reflecting the position of (*T, p, q*) within *T*. Similarly, *j* = *j*^*′*^ + *q* − 1 holds, and hence (*T, p, q*) is recovered by (*str*(*S, i*^*′*^, *j*^*′*^), *i* − *i*^*′*^ + 1, *j* − *j*^*′*^ + 1).

We need (*S, i*^*′*^, *j*^*′*^) ∈ *ProjOcc*_*𝒰*_ to finish showing the second statement. If *T* is in 𝒰, then trivially (*S, i*^*′*^, *j*^*′*^) = (*S*, 1, |*S*|) holds and (*S*, 1, |*S*|) ∈ *ProjOcc*_*𝒰*_ is applied by item 1. in Definition S11. Otherwise, *T* is in *R*_𝒰_, then (*S, i*^*′*^, *j*^*′*^) is *occ*_*𝒰*_ (*T*)_**1**_ because the first occurrence of some subtring co-occurring within *T* was utilized for identifying (*S, i, j*). Proposition 5 confirms that *occ*_*𝒰*_ (*T*)_**1**_ ∈ *ProjOcc*_𝒰_.

Lastly, assume some (*S, x, y*) in *ProjOcc*_𝒰_ satisfies (*S, i, j*) ⊊ (*S, x, y*) ⊊ (*S, i*^*′*^, *j*^*′*^). Then a maximal repeat *R*^*′*^(:= *str*(*S, x, y*)) exists such that *R*^*′*^ is a superstring of *R*(:= *str*(*T, p, q*)) and *T* is a superstring of *R*^*′*^. Then *R*^*′*^ could be captured by extending (*T, p, q*) and *R*^*′*^ occurs more than *T*, so (*T, p, q*) is not a locally-maximal repeat region, leading to a contradiction.

For the backward direction, assume (*T, p, q*) is not a locally-maximal repeat region for contradiction. Then some locally-maximal repeat region (*T, p*^*′*^, *q*^*′*^) must be found by extending (*T, p, q*) because *R* := *str*(*T, p, q*) occurs more than *T*. Then by following the the same steps in the first paragraph utilizing Theorem S6 and Definition S11, (*T, p*^*′*^, *q*^*′*^) is used to identify an occurrence (*S*^*′′*^, *i*^*′′*^, *j*^*′′*^) in *ProjOcc*_𝒰_ that shares some letter extension. So, letting *R*^*′*^ := (*T, p*^*′*^, *q*^*′*^), a substring of form either *σ*_*l*_*R*^*′*^ or *R*^*′*^*σ*_*r*_ is used to find a first occurrence and the substring co-occurs within *T*, so (*S*^*′′*^, *x*^*′′*^, *y*^*′′*^) appears within *occ*_𝒰_ (*T*)_**1**_ so that its relative occurrence within *occ*_𝒰_ (*T*)_**1**_ is (*T, p*^*′*^, *q*^*′*^). Therefore, (*S*^*′′*^, *i*^*′′*^, *j*^*′′*^) ⊊ *occ*_𝒰_ (*T*)_**1**_. Also, (*S, i, j*) ⊊ (*S*^*′′*^, *i*^*′′*^, *j*^*′′*^) because (*T, p*^*′*^, *q*^*′*^) is an extension of (*T, p, q*) that (*S*^*′′*^, *i*^*′′*^, *j*^*′′*^) is an extension of (*S, i, j*) too. Lastly, *occ*_𝒰_ (*T*)_**1**_ = (*S, i*^*′*^, *j*^*′*^) indeed holds because of (*S, i, j*) ⊊ (*S, i*^*′*^, *j*^*′*^). That is, if *occ*_𝒰_ (*T*)_**1**_ is not (*S, i*^*′*^, *j*^*′*^) but some other occurrence of *T* in *S*^*′′*^, then the corresponding occurrence of *R* in *ProjOcc*_𝒰_ must be found in *S*^*′′*^ instead of (*S, i, j*). Thus, the presence of (*S*^*′′*^, *i*^*′′*^, *j*^*′′*^) leads to contradiction. □

Therefore, two occurrences in their “closest” containment relationship in *ProjOcc*_𝒰_ imply their strings form a locally-maximal repeat region. Since every occurrence in *ProjOcc*_𝒰_ originates from either a maximal repeat or a sequence, each represents a vertex in the Pro*k*rustean graph. Therefore, collecting these relative occurrences constructs the edge set of the Pro*k*rustean graph. Refer to Section S3 for the algorithm.

#### S3.3 Implementation

We briefly introduce the techniques implemented in our code base: https://github.com/KoslickiLab/prokrustean.

Firstly, the BWT is constructed from a set of sequences using any modern algorithm supporting multiple sequences [17, 21, 30]. The resulting BWT is then converted into a succinct string representation, such as a wavelet tree, to facilitate access to the “nodes” of the implied suffix tree. For this purpose, we used the implementation from the SDSL project [24]. The traversal is built on foundational works by Belazzougui et al. [4] and Beller et al. [6], as detailed by Nicola Prezza et al. [37]. Their node representation, based on LCP intervals, enables constant-time access to *occ*_*𝒰*_ (*Rσ*) and *occ*_*𝒰*_ (*σR*) for each substring *R* and letter *σ* ∈ *Σ*. This capability is crucial for verifying the maximality conditions outlined in Definition S11. Also, the first occurrences *occ*_*𝒰*_ (*Rσ*)_**1**_ and *occ*_*𝒰*_ (*σR*)_**1**_ can be identified as the start of the LCP interval of *Rσ* and *σR* within the representation, respectively.

Therefore, *ProjOcc*_*𝒰*_ is collected by exploring the nodes of the suffix tree implicitly supported by the BWT of *𝒰*. The exploration requires *O*(log |*Σ*| *·* |*T*_𝒰_ |) time in total, because succinct string operations take *O*(log |*Σ*|) time to move over and extract node representations. Identifying a maximal repeat takes *O*(|*Σ*|) time and then collecting its first occurrences used in *ProjOcc*_𝒰_ takes *O*(|*Σ*|^2^), because each combination of occurrence, e.g., |*occ*_𝒰_ (*σ*_*l*_*Rσ*_*r*_)|) is accessible in constant time. In summary, constructing *ProjOcc*_𝒰_ takes *O*(log |*Σ*| *·* |*Σ*|^2^ *·* |*T*_*𝒰*_ |) time and *O*(|*BWT* | + |*ProjOcc*_𝒰_ |) space where |*BWT* | is the size of the succinct string representing the BWT of 𝒰.

Lastly, reconstructing the Pro*k*rustean graph using Theorem S7 takes *O*(|*ProjOcc*_𝒰_ |) time, which is introduced in Section S3. Note that the computation need not consider every combination of occurrences in *ProjOcc*_𝒰_ ; it leverages the property that an occurrence (*S, i, j*) ∈ *ProjOcc*_𝒰_ can imply locally-maximal repeat regions with up to two extensions in *S* found in *ProjOcc*_𝒰_. Therefore, *ProjOcc*_𝒰_ is grouped by each sequence *S* ∈ 𝒰, and the edge set *E*_𝒰_ can be built from each group. Hence, the total space usage is *O*(|*BWT* | + |*ProjOcc*_𝒰_ | + |𝒢_𝒰_ |), where *O*(|*ProjOcc*_𝒰_ |) = *O*(|𝒢_𝒰_ |). The space usage is primarily output-dependent that around 2|𝒢_𝒰_ | in practice, and the most time-consuming part is the construction of *ProjOcc*_𝒰_ via traversing the implicit suffix tree.

#### S3.4 Algorithm: from projected occurrences to Pro*k*rustean graph

The algorithm below reduces the task of reconstructing the Pro*k*rustean graph from *ProjOcc*_*𝒰*_ to a problem designed for a specific string *S*∈ 𝒰. For the subset of regions {(*S*^*′*^, *i, j*) ∈*ProjOcc*_𝒰_ |*S*^*′*^ = *S*}, determine their inclusion relationships to identify the closest related pairs. This is easily parallelized as it processes the intervals for each *S* separately. And the running time is linear to the size of the subset of regions per each *S*, hence |*ProjOcc*_𝒰_ | in total.

An important assumption in the algorithm is that for any region (*S, i, j*) ∈ *ProjOcc*_𝒰_, within the subset {(*S*^*′*^, *i, j*) ∈*ProjOcc*_𝒰_ |*S*^*′*^ = *S*}, there are at most two regions (*S, i*^*′*^, *j*^*′*^) such that (*S, i, j*) ⊊ (*S, i*^*′*^, *j*^*′*^) and no (*S, x, y*) exists where (*S, i, j*) ⊊ (*S, x, y*) ⊊ (*S, i*^*′*^, *j*^*′*^). It is straightforward to check the property.

##### Algorithm S5 Find recursive inclusions of intervals

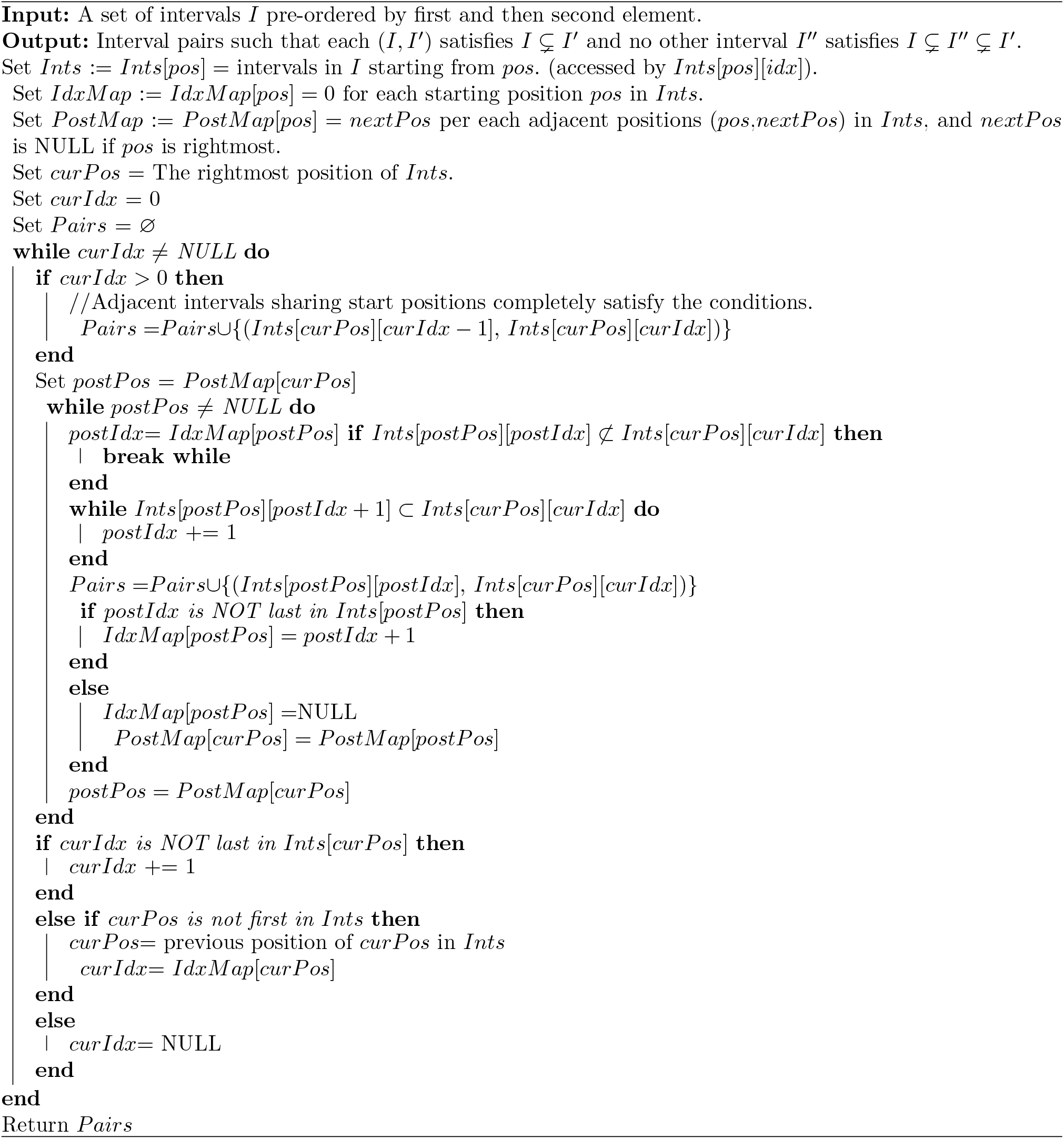

## Footnotes

1 https://github.com/KoslickiLab/prokrustean

